# Dynamics of the adhesion complex of the human pathogens *Mycoplasma pneumoniae and Mycoplasma genitalium*

**DOI:** 10.1101/2023.07.31.551205

**Authors:** David Vizarraga, Akihiro Kawamoto, Marina Marcos-Silva, Jesús Martín, Fumiaki Makino, Tomoko Miyata, Jorge Roel-Touris, Enrique Marcos, Òscar Q. Pich, David Aparicio, Ignacio Fita, Makoto Miyata, Jaume Piñol, Keiichi Namba, Tsuyoshi Kenri

## Abstract

*Mycoplasma pneumoniae* is a bacterial wall-less human pathogen and the etiological agent of atypical pneumonia and tracheobronchitis in both adults and children. *M. pneumoniae* infectivity, gliding motility and adherence to host target respiratory epithelial cells are mediated by adhesin proteins P1 and P40/P90 forming a transmembrane complex that binds to sialylated oligosaccharides human cell ligands. Here we report the cryo-EM structure of P1 bound to the Fab fragment of monoclonal antibody P1/MCA4, which stops gliding and induces detachment of motile *M. pneumoniae* cells. On the contrary, polyclonal antibodies generated against the N-domain of P1 or against the whole ectodomain of P40/P90 have little or no effects on adhesion or motility. The epitope of P1/MCA4, centred on loop Thr1426-Asp1438 in the small C-terminal domain of P1, is inaccessible to antibodies in the “open” conformation of the adhesion complex, when ready for attachment to sialylated oligosaccharides. Mutations in the highly conserved Engelman motifs found in the transmembrane helix of P40/P90 also alter adhesion and motility. During the attachment/detachment cycle of the adhesion complex, the C-terminal domain of P1 experiences large conformational rearrangements that are hindered by the antibodies against the domain. Interfering with the gliding of mycoplasma cells suggests new ways to confront *M. pneumoniae* infections.

## INTRODUCTION

*Mycoplasma pneumoniae* (*Mycoplasmoides pneumoniae*) is one of the leading microorganisms causing community acquired bacterial pneumonias ^1^. Coinfections of *M. pneumoniae* with other respiratory pathogens, such as SARS-CoV-2, Adenovirus or *Chlamydia pneumoniae*, have been associated with importantly increased morbidities ^1–3^. Recent pneumonia outbreaks in Europe and Asia due to the re-emergence of *M. pneumoniae* also underlines the clinical relevance of this human pathogen ^4^. However, the tropism for respiratory tissues of *M. pneumoniae* might be exploited for biomedical applications, and attenuated strains of this mycoplasma have been recently engineered to be used as living pills to treat pulmonary diseases ^5^.

*M. pneumoniae* exhibits a membrane protrusion at one of cell poles, referred as the “attachment” or “terminal” organelle (TO), that is instrumental for infection and gliding motility. *M. pneumoniae* cells bind to glass surfaces coated with sialylated oligosaccharides (SOs), which are abundant on the human epithelial surfaces, and glide in the direction of the TO ^6–8^. The type of motility presented by *M. pneumoniae* is shared only by mycoplasma species belonging to the pneumoniae cluster ^7,9^. The molecular machinery for motility is located in the TO and involves about fifteen different proteins organized as surface structures and internal parts ^9–11^. The main and most abundant surface structure is the adhesion complex, also known as the Nap complex, which comprises the transmembrane adhesin proteins P40/P90 and P1 ^12–14^. A similar organization of the adhesins P110 and P140 forming a 540 kDa membrane complex is found in the closely related human pathogen *Mycoplasma genitalium* ^11,15^. Adhesins play an essential role in attachment to surfaces by binding to the SOs ligands ^15,16^.

During the last few years, several high-resolution structures have been determined for adhesins P1 and P40/P90 ^14^ and also for the closely related orthologous from *M. genitalium* P140 and P110 ^17–19^. The Nap complexes were proposed to cycle between “open” and “closed” conformations where the SOs binding pockets become accessible or inaccessible, respectively. The SO binding pocket is located in the P40/P90 N-terminal domain, although accessibility to the pocket is determined by the interaction with the N-terminal domain of P1 that is tighter in the “closed” conformation. Adhesins P1 and P40/P90, together with protein P116, are the most immunogenic proteins of *M. pneumoniae* ^14,20^. To study the immunogenic and functional properties of P1, monoclonal antibodies were generated against a fragment of this adhesin spanning from residue Ala1160 to Gln1518, (P1:1160-1518 peptide) (**Supplementary Table I**). One of the monoclonal antibodies obtained (henceforth referred as P1/MCA3) was reported some years ago to progressively reduce gliding speed, removing mycoplasma cells from surfaces in a concentration-dependent way, while having little effect on non-moving cells ^21^. These observations led to the conclusion that during gliding, P1 has to experience important conformational changes, participating like a leg in a “power stroke” that propels the cell, with epitope exposure to antibody P1/MCA3 significantly reduced in non-moving cells.

How the functioning of the Nap complex is related to the motility of mycoplasma cells remains elusive. Monoclonal antibody P1/MCA3 is no longer available, but we have now solved by cryo-electron microscopy (cryo-EM) the structure of P1 in complex with the Fab fragment of P1/MCA4, which was obtained at the same time as P1/MCA3 against the same P1:1160-1518 peptide (**Supplementary Table I**). P1/MCA4 also affects motility and enhances detachment of moving mycoplasma cells similarly to P1/MCA3. The determined structure provides a snapshot of the “closed” conformation of P1 that prevents the Nap complex from attachment to SOs. We also investigated how motility is affected by polyclonal antibodies against the whole ectodomains of adhesins P1 and P40/P90 and those against constructs of the N-terminal domain of P1. Mutations in the highly conserved Engelman motifs, found in the transmembrane helices of adhesins, can also alter adhesion and motility, providing information about how extra- and intra-cellular regions communicate with each other. Altogether, these results provide a deep insight into the Nap complex function and explain the specific neutralization mechanism deployed by antibodies against the C-terminal domain of P1.

## RESULTS

### Monoclonal antibody P1/MCA4 and its binding to P1

Monoclonal antibody P1/MCA4, raised against the P1:1160-1518 peptide, exhibited strong hemadsorption inhibitory activity for *M. pneumoniae.* In the presence of P1/MCA4, gliding mycoplasma cells reduce rapidly their speed and eventually halt and detach from sialylated glass surfaces (**Video I**), while non-moving cells are not affected by the presence of P1/MCA4 and remain attached to surfaces. As mentioned in the introduction, a similar behaviour had been reported for antibody P1/MCA3 ^21^. It is worth mentioning that this behaviour is also observed in three additional monoclonal antibodies raised in the same batch as P1/MCA3 and P1/MCA4 (**Supplementary Table I**). The P1:1160-1518 peptide corresponds to a fragment of the N-terminal domain (residues Ala1160-Thr1399) and to (almost) the whole C-terminal domain (Ala1400-Gln1518) according to the structural information now available ^14^.

cDNA sequencing of P1/MCA4 mRNAs in hybridoma cells showed the presence of a single IgG heavy chain mRNA but two different light chain mRNAs (**Supplementary** Figure 1). N-terminus analysis of samples from antibody P1/MCA4 (obtained as indicated in the Material and Methods) confirmed the presence of one heavy chain but two different light chains (roughly 70% for the most abundant). The heterogeneity of light chains might explain the failure of numerous crystallization attempts involving P1/MCA4. Analysis by multi-angle light scattering (MALS) showed the formation of a stable and homogeneous P1-Fab(P1/MCA4) complex, in a solution with equimolecular amounts of P1/MCA4 Fabs and the whole P1ectodomain (residues Thr29-Asp1521), indicating that the two kinds of Fab found in P1/MCA4 bind efficiently to P1 (**Supplementary** Figure 2).

Antibody P1/MCA4 presents high affinity for a construct corresponding to the C-terminal domain of P1, proving that this small domain contains the epitope. Moreover, western blotting analysis performed using different segments of P1 showed that residues Thr1426-Asp1438 are a key part of the epitope (**Supplementary** Figure 3). An additional observation that could have clinical implications is that P1/MCA4 presents affinity also for the C-terminal domain of P140 (residues Lys1220-Asp1351) from *M. genitalium,* even stopping the movement of cells, although after longer treatments than with *M. pneumoniae*, in agreement with the high sequence and structural similarities between the C-terminal domains of the adhesins of *M. pneumoniae* and *M. genitalium* (**Supplementary** Figure 4).

### Cryo-EM structure of the P1-Fab(P1/MCA4) complex

The structure of the ectodomain of P1 bound to the Fab fragment of P1/MCA4 was determined by cryo-EM single particle analysis with an overall resolution in the final map of 2.4 Å (**Figure 1a, Supplementary** Figures 5-6 and **Supplementary Table II**). The high quality of the map allowed accurate modelling of the structure from P1 and the Fab variable domains, defining clearly the paratope-epitope interface. The Fab constant domains were less well defined, indicating some flexibility in the Fab elbow. The sequence observed for the Fab corresponded to the one with light chain L1. The structure of P1 consists of a large N-terminal domain (residues 60-1399) and a smaller C-terminal domain (1400-1521), in full agreement with the structures of P1 reported previously ^14^. Superposing the P1 structure determined here with the P1 crystal structure (PDB code 6rc9), gives a root mean square deviation (rmsd) of 1.80 Å for 1236 aligned residues (from a total of 1339 in the model) (**Supplementary** Figure 7). When the superposition is done separately for the N-terminal and C-terminal domains, rmsds are 1.65Å (1151 aligned residues out of 1226) and 1.20Å (97 aligned out of 104), respectively. These results show the high plasticity of P1, in particular for the N-terminal domain, which experiences significant changes although it participates only indirectly in the interaction with the P1/MCA4 Fab. Comparison of the crystal (6rc9) and the new (in the complex of P1-Fab) structures of P1, indicates a 7.7° rotation of the C-terminal domain relative to the N-terminal domain, confirming hinge movements between the two domains as had suggested the cryo-EM structure of P1 where the C-terminal domain was disordered ^14^. The structure of the P1-Fab complex shows that the epitope of P1/MCA4 is conformational, although involving residues from the C-terminal domain only, in particular from loop Val1425-Asp1438 in agreement with the epitope mapping analysis (**Figure 1a and Supplementary** Figure 3). The quality of the cryo-EM map was high enough to even allow the localization in the N-terminal domain of a significant number of solvent molecules, most of them also present in the crystal structure of P1 (**Supplementary** Figure 6). It is worth mentioning that docking using a previous cryo-EM map determined at lower resolution, correctly predicted the overall binding interface between P1 and the Fab(P1/MCA4), with the largest deviations arising from the inaccuracy on the fitting of the P1 C-terminal domain (**Supplementary** Figure 8).

**Figure 1.**
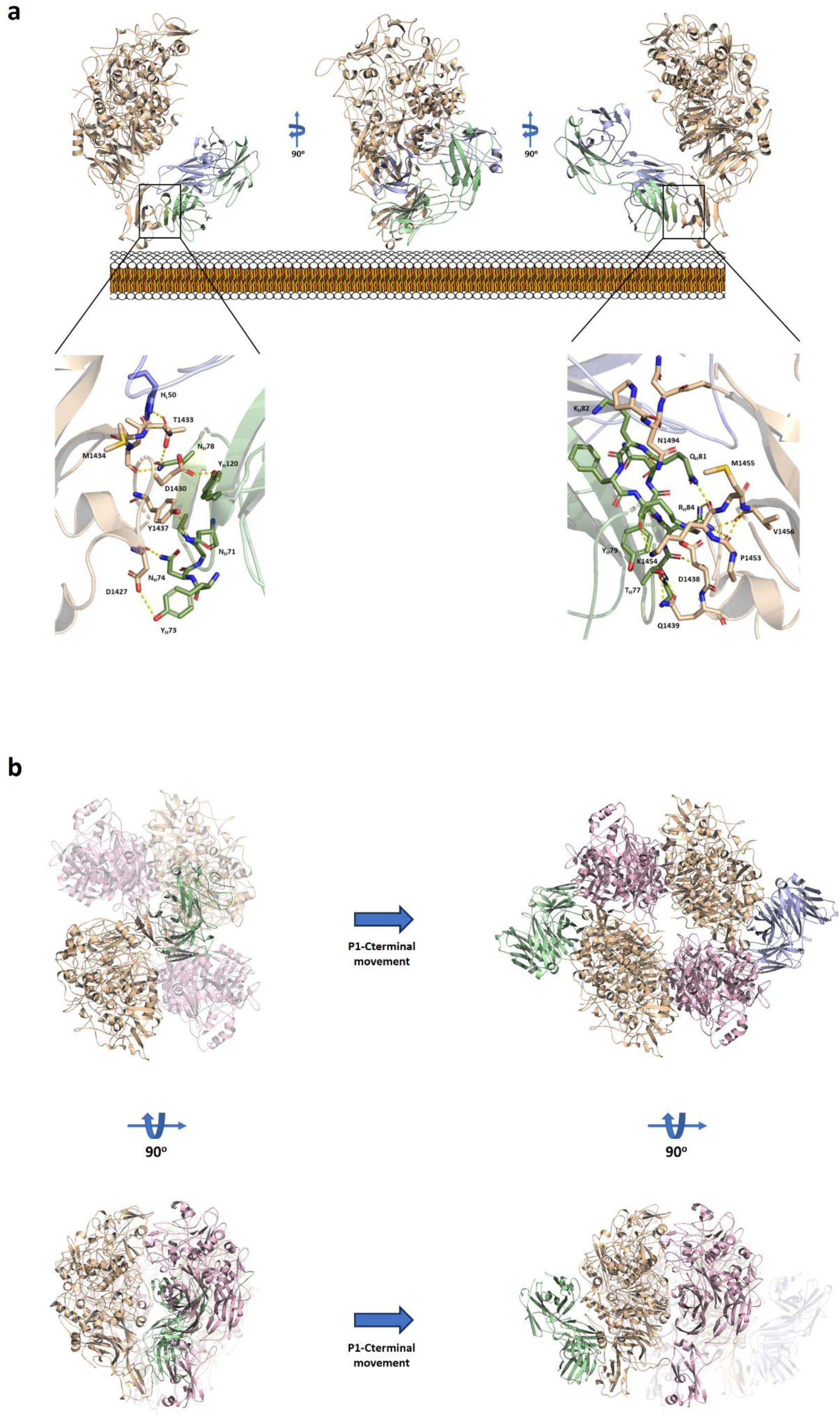
Cryo-EM structure of the P1-Fab(P1/MCA4) complex. **a)** Ribbon representations, with three 90° apart views, of the complex between P1 (brown) and the Fab fragment of monoclonal antibody P1/MCA4 (light and heavy chains in blue and green, respectively). Insets showing the recognition interactions between P1 and P1/MCA4. The epitope is located in the C-terminal domain of P1, close to the mycoplasma membrane displayed as a reference. **b)** Two 90° apart views of the Nap complex in the “open” conformation with one of the P1 subunits replaced by the P1-Fab(P1/MCA4) complex (left panels). The P1/MCA4 epitope, buried at the center, is totally inaccessible to antibodies in the “open” conformation and the Fab would catastrophically clash with the adhesins. The C-terminal domain has to experience an important hinge rearrangement to expose the epitope avoiding steric clashes (right panels).

Importantly, superposing the structure of P1-Fab(P1/MCA4) onto the Nap complex in “open” conformation available for *M. genitalium* ^17^ results in a catastrophic steric clash of the Fab against the adhesins, indicating that the epitope of P1/MCA4 is totally inaccessible to antibodies in the “open” conformation of the Nap complex (**Figure 1b**). Therefore, this complex must experience important structural rearrangements between the “open” conformation and when the epitope of P1/MCA4 is fully exposed.

### Attempts to obtain the ternary complex P40/P90-P1-Fab(P1/MCA4)

A solution with equimolecular amounts of Fab(P1/MCA4) and of ectodomains from P1 (residues 29-1521) and from P40/P90 (residues 23-1114) produced a mixture of several complexes where the ternary complex P40/P90-P1-Fab(P1/MCA4) was not clearly identified, while heterodimers P40/P90-P1 and P1-Fab(P1/MCA4) were abundant (**Supplementary** Figure 9). Cryo-EM confirmed the mixture of different complexes in the classifications **(Supplementary** Figure 10). Attempts to determine the structure of a ternary complex gave a map with an overall resolution lower than 10Å **(Supplementary Table II)**. The best-defined part of this map corresponds to the N-terminal domains of P40/P90 and P1 that interact tightly with each other similarly to what had been determined for the “closed” structures of orthologue adhesins P110 and P140 from *M. genitalium* ^17^. Density corresponding to the C-terminal domain of P1 was quite strong, although fitting of the domain presented some ambiguities due to the limited resolution and small size of the domain. Density was missing for the C-terminal domain of P40/P90 and uninterpretable for the Fab. Docking the structure of the P1-Fab binary complex by superimposing the N-terminal domains of P1, resulted in steric clashes between the Fab and P40/P90 although their footprints on P1 do not overlap, suggesting that P40/P90 might compete with the Fab for the formation of the ternary complex **(Supplementary** Figure 11). However, steric problems could be relieved by a hinge rotation of the P1 C-terminal domain and the scarcity of the ternary complex shows that the energy required by the hinge rotation is larger than the energy provided by the binding to P1 of either P40/P90 or the Fab.

### Polyclonal antibodies against the P40/P90 and P1 ectodomains and against the N-terminal domain of P1

Polyclonal antisera against the ectodomains of P40/P90 (residues Ala25-Pro1113) and of P1(residues Thr29-Asp1521) were obtained by immunizing mice with recombinant versions of these polypeptides (**Table I** and **Supplementary Table III**). Polyclonal antisera against the N-terminal domain of P1 were obtained by immunizing mice with the P1 N-terminal domain (residues Thr29-Ala1375). These antisera were tested by ELISA, Western Blotting and immunofluorescence assays on whole *M. pneumoniae* cells, and titers higher than 1/2000 were obtained for all the antisera. Effects of these antisera on *M. pneumoniae* motility and attachment were investigated by incubating *M. pneumoniae* cells in the presence of 3% gelatin in the SP4 medium ^16^. Because *M. pneumoniae* cells spontaneously detach at high frequencies from the observation surface in the absence of gelatin ^20^ more reproducible and consistent results were obtained when gliding motility was analyzed in SP4 medium supplemented with gelatin. In the absence of antibodies (negative control), most of the cells remained attached and motile during the observation time (**Table I)**. In contrast, the P1 ectodomain antisera stopped the movement of 50% of mycoplasma cells after four minutes of incubation (**Table I, Video II)** with very few cells remaining motile at the end of the observation period. No significant effects on mycoplasma gliding were detected neither for the antisera against the P1 N-terminal domain nor for the P40/P90 ectodomain, even at the lowest dilution tested **(Table I, Videos III and IV)**. It is also worth mentioning that cell adhesion was inhibited when cells were added to media that already contained antisera against the ectodomains of either P1 or P40/P90, suggesting that these antibodies can interfere with their binding to SOs via steric hindrance, but their epitopes are not accessible when the cells are attached to surfaces.

**Table I.**

| Antibody<br>or sera | SP4 |  |  |  | SP4 3% gelatin |  |  |
| --- | --- | --- | --- | --- | --- | --- | --- |
|  | Before # | After # |  |  | Before # | After # |  |
| | Motile cells<br>(% ± SE) | Motile cells<br>(% ± SE) | RT<sub>50</sub> (min<br>± SE) & | Detached cells<br>(% ± SE) | Motile cells<br>(% ± SE) | Motile cells<br>(% ± SE) | RT<sub>50</sub> (min<br>± SE) & |
| **Control** § | 89.9 ± 1.2 | 76.1 ± 4.8 | NA | 51.5 ± 4.3 | 96.3 ± 0.3 | 96.5 ± 0.1 | NA |
| **P1** | 89.4 ± 0.6 | 1.8 ± 1.8 | 4.65 ± 1.2 | 72.3 ± 2.4 | 96.5 ± 0.5 | 3.4 ± 2.8 | 4.1 ± 0.35 |
| **P40/P90** | 87.5 ± 3.9 | 74.3 ± 4.4 | NA | 45.2 ± 26.1 | 96.4 ± 0.4 | 95.7 ± 0.3 | NA |
| **P1 N-ter** | 95.6 ± 0.1 | 85.4 ± 1.6 | NA | 59.8 ± 0.1 | 88.9 ± 3.2 | 88.2 ± 2.7 | NA |
| **P1/MCA4** | - | - | - | - | 97.3 ± 0.7 | 0 ± 0 | 2.6 ± 0.26 |

§ Negative control. PBS with no antibody was added.

Binding of antibodies to epitopes whose accessibility varies during a conformational transformation interferes with the transformation. According to this idea, the attachment/detachment cycle is halted by antibody P1/MCA4 against the C-terminal domain of P1 that experiences important rearrangements during the cycle. Instead, the attachment/detachment cycle is not altered, or very little, by antibodies against P40/P90 or against the N-terminal domain of P1, indicating that accessibility of their epitopes must remain essentially unchanged during the cycle.

### Mutations in the Engelman motifs

The transmembrane domains of P1 and P40/P90, predicted to be single helices, exist just after the C-terminal domains of these proteins. Remarkably, these helices contain one Engelman motif in P1 and two Engelman motifs in P40/P90 (**Supplementary** Figure 12). These three Engelman motifs are highly conserved in the pneumonia cluster of mycoplasmas. Engelman motifs, frequently involved in high-affinity interactions between membrane helices, have the sequence GXXXG, with X being any residue but in general hydrophobic (Russ & Engelman, 2000). We have investigated by mutational analysis, possible contributions of the Engelman motifs from adhesins to the organization and functioning of the Nap complex. The study was performed with *M. genitalium* because simple genetic methods to introduce point mutations are available in this microorganism and transmembrane helices from orthologue adhesins of *M. pneumoniae* and *M. genitalium* present high sequence identities (**Supplementary** Figure 12). The Engelman motif at the C-end of the P140 transmembrane helix, close to the internal side of the cell membrane, will be referred as E1 and the two Engelman motifs in P110 as E2 and E3, close to the external and to the internal sides of the cell membrane, respectively (**Figure 2a**). Mutations were introduced by transposon delivery in a *M. genitalium* null mutant G37ΔAdh (G37ΔMG_191-ΔMG_192), to obtain isogenic strains bearing Gly to Phe double point mutations of the glycine residues of P140 and/or P110 Engelman motifs (**Supplementary Table IV**). The protein profile analysis of these mutant strains showed that adhesins bearing the different mutations were expressed at similar levels to P140 and P110 in wild type (WT) cells (**Supplementary** Figure 13). The properties of the adhesins variants were tested by quantitative hemadsorption assay (**Figure 2b**). The E1 mutant strain cells showed hemadsorption values very similar to those of WT cells and also similar to what is obtained when a WT copy of both adhesin coding genes were delivered by transposition to the null mutant G37ΔAdh. In contrast, E2 mutants showed an intermediate adherence phenotype and E3 mutants had a more severely impaired hemadsorption phenotypes, characterized by low B_max_ values (see in Material and Methods). As expected, E1-E2 and E1-E3 mutant strains exhibited phenotypes similar to E2 and E3, respectively. In addition, E2-E3 and E1-E2-E3 mutant strains showed a non-adherent phenotype characterized by very low B_max_ and very high K_d_ values, similar to the ones of the G37ΔAdh null mutant strain. Immunolabeling with a polyclonal antiserum against the Nap complex from *M. genitalium*, seemed to indicate that labelling concentrates at the tip of the triple mutated (E1-E2-E3) cells, as found for the P110 and P140 WT adhesins (**Supplementary** Figure 14). Cells from E1 mutants exhibited normal morphologies when analyzed by Scanning Electron Microscopy. Remarkably, all cell variants where E2 or E3 was mutated showed a multiple TO phenotype. The fraction of cells presenting multiple TOs (MOTs) increased from a small percentage in E2 variants to most cells in E2-E3 and E1-E2-E3 variants (**Figure 2c**). Nascent TOs develop adjacent to a preexistent TO with gliding providing the strength to deliver the new TOs to the opposite cell pole, a process that is hindered in cells having adhesion and motility deficiencies ^22^, which can explain the correlation found in the Engelman mutated variants between reduced adhesion and increasing rates of MOT cells. Moreover, it had been shown that P110 and P140 adhesins are essential for TO development ^23^, which indicates that these adhesins are properly folded and positioned in the TOs of the Engelman variant mutant cellss. Therefore, mutational analysis allows to conclude that Engelman motifs from P110, but unexpectedly not from P140, have a critical and synergic contribution to adhesion, strongly suggesting that interactions between the transmembrane helices of the two P110 subunits are required to achieve a functional “open”, ready for binding, conformation of the Nap complex.

**Figure 2.**
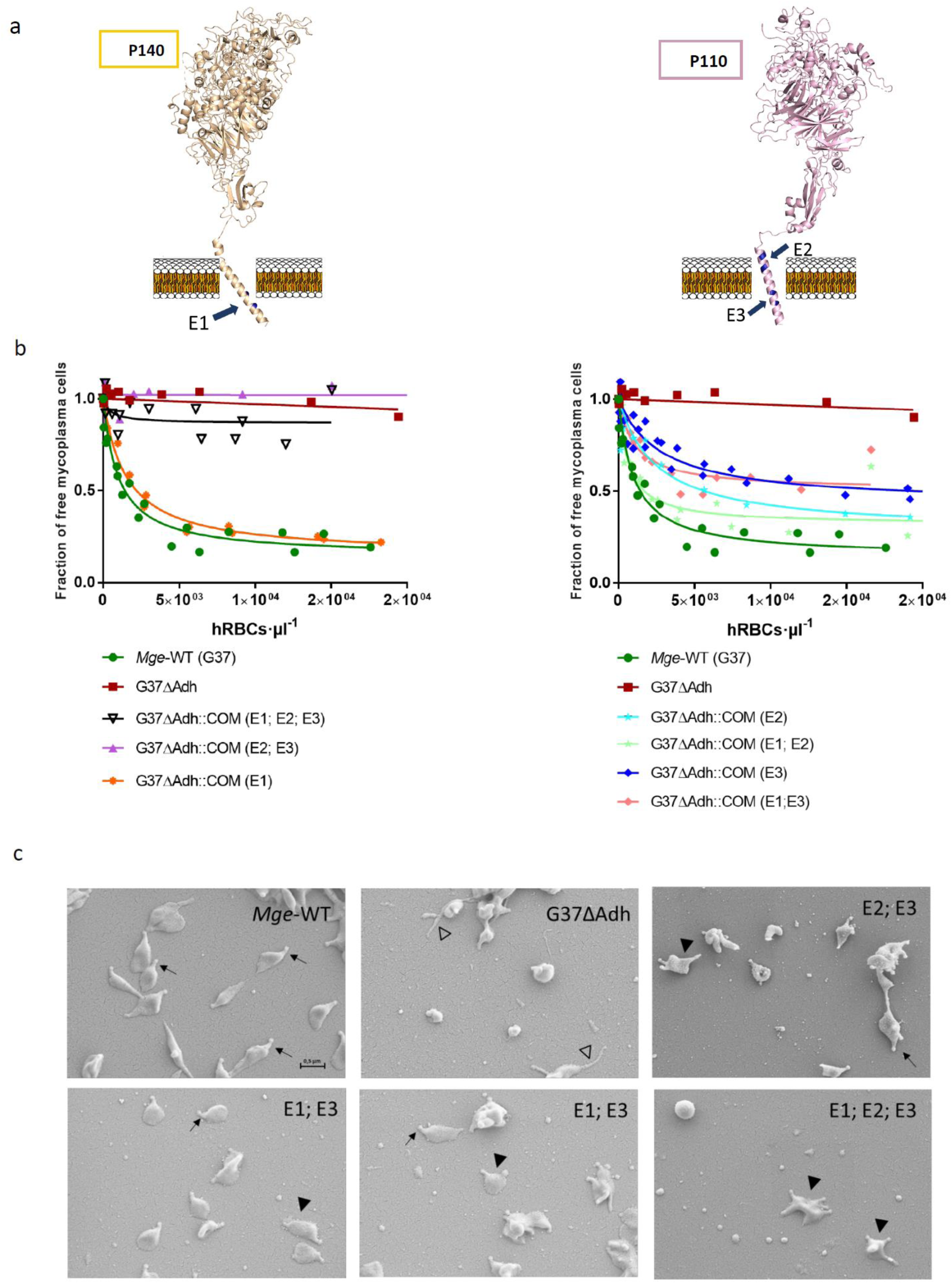
Engelman motifs in adhesins. **a)** Transmembrane helices of adhesins from the pneumonia cluster of mycoplasmas contain three highly conserved Engelman motifs (GxxxG sequences), referred as E1 (P140) and E2, E3 (P110) in *M. genitalium*. **b)** Mutational analysis, performed in the Engelman motifs of P140 and P110, indicate that motifs E2 and E3 from P110 have important and synergic contributions to adhesion. On the contrary, no effects were observed when E1 from P140 was mutated. **c)** Changes in adhesion correlate with increasing rates of cells presenting multiple TOs phenotypes.

## DISCUSSION

The pneumonia cluster of mycoplasmas has developed a unique, strikingly complex, molecular machinery for gliding motility that consists of internal and surface structures located in the TO, a morphologically conspicuous cell protrusion. A gliding model has been proposed based mainly on biophysical and structural data of the internal structures (Kawamoto et al., 2016; ^7,9^. The model is a variant of the “inchworm” model, in which repeated contractions and extensions, synchronized with alternative attachment and detachment of the TO front and rear sides, enable the smooth gliding of cells ^6,24,25^. According to this model, adhesins P1 and P40/P90 forming the Nap complex, which is the main surface structure of the TO, hve a key contribution to both attachment and movement, by going throughout an iterative four stages cycle during gliding (Mizutani et al., 2021). In the last few years a wealth of high resolution structural data has been obtained for the Nap complex ^11,15^.

The structure of the complex of *M. pneumoniae* adhesin P1 and the Fab fragment from the monoclonal antibody P1/MCA4, determined in this work, shows that the P1/MCA4 epitope involves residues of the C-terminal domain of P1 only **(Figure 1a)**. This epitope is totally inaccessible to antibodies in the “open” (ready for binding to SOs) conformation of the Nap complex **(Figure 1b)**. Therefore, for the P1/MCA4 epitope to be exposed, the adhesion complex has to experience a major rearrangement with respect to the “open” conformation. The conformation where P1/MCA4 binds must be unsuitable for attachment to SOs, as binding of antibody P1/MCA4 slows and induces detachment of moving *M. pneumoniae* cells only. Together with the available information, our results allow now to model accurately the four stages of the Nap complex cycle (**Figure 3, Video V**) explaining how internal and surface structures interact with each other.

**Figure 3.**
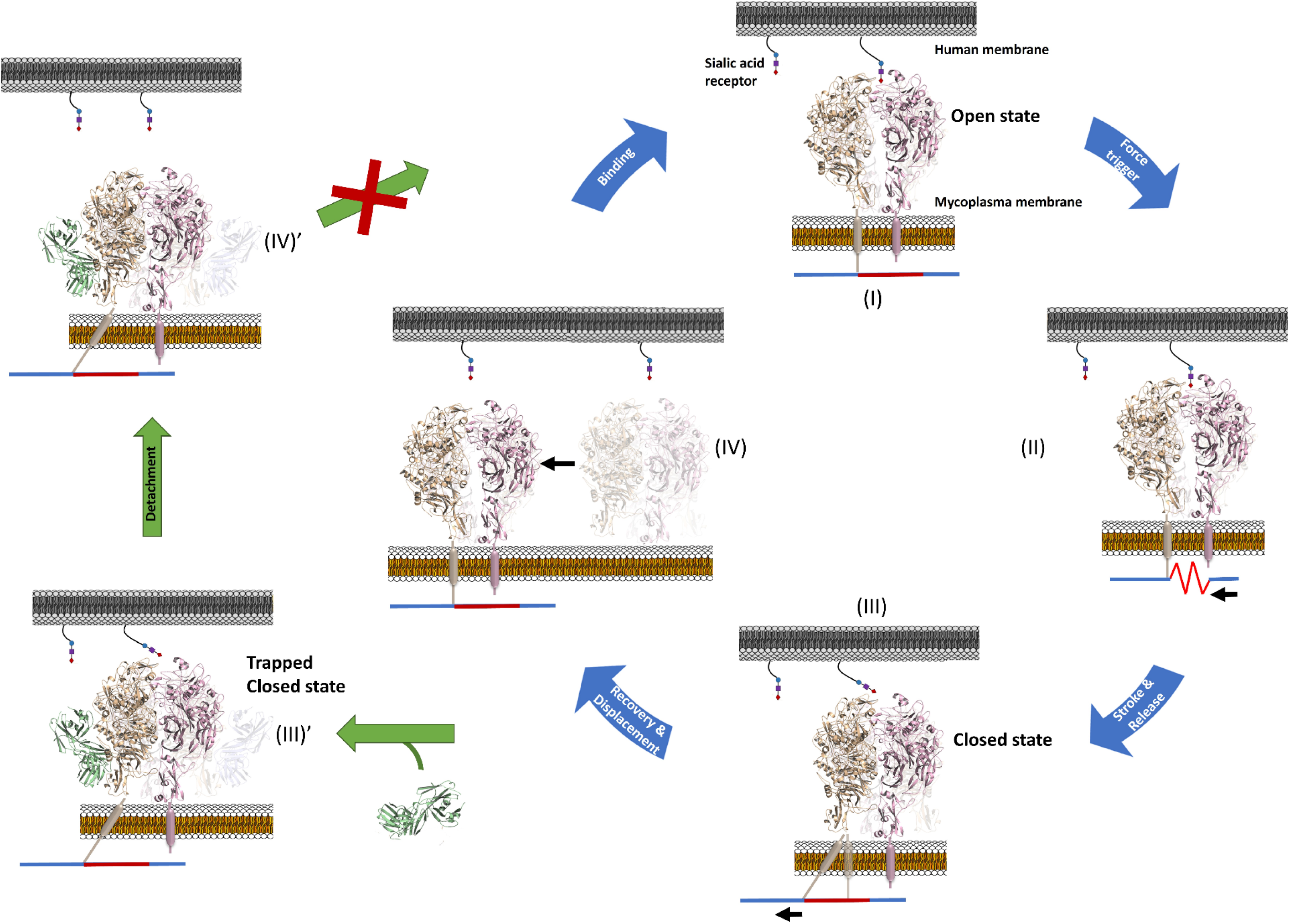
The attachment and detachment cycle of the Nap complex. The Nap complexes participating in adhesion and motility cycle between “open” and “closed” states, with the binding site to sialic oligosaccharides becoming alternatively accessible and inaccessible for binding. During the cycle, structural rearrangements are experienced mainly by the C-terminal domain of P1, with hinge rotations of about 175°. Movements of the C-terminal domain of P1 may be driven by the force, generated by ATP hydrolysis, transmitted from the back end of the TO (at the right of images) through the rod, which is represented as a line composed of elastic (red) and rigid (blue) parts. Displacements are shown by black arrows. Binding of the monoclonal antibody P1/MCA to the C-terminal domain of P1 halts the cycle, trapping the “closed” conformation and inducing detachment in motile cells only.

Stage I: The cycle starts with the “open” conformation that is ready for binding to the SOs. The extracellular region from the “open” conformation of the Nap complex was built according to the structure determined *in situ* by cryo-electron tomography (cryo-ET) for *M. genitalium* ^17^. The Engelman motifs found in the transmembrane helix of P40/P90 have a critical role to stabilize the “open” conformation, suggesting that the helices from the two P40/P90 subunits interact with each other.

Stage II: Binding to SOs triggers the transition of the Nap complex towards the “closed” conformation. During the transition the exposed surfaces of the P40/P90 subunits remain unchanged, in agreement with the restraints imposed by the interactions between the transmembrane helices of the two P40/P90 subunits and previous studies showing that the two SOs binding sites of the Nap complexes are probably involved ^26,27^. In turn, the C-terminal domain of P1 has to experience an important rearrangement to allow the exposure of the P1/MCA4 epitope (**Supplementary** Figure 15).

Stage III: Reaching the “closed” conformation of the Nap complex, where the N-terminal domains of P1 and P40/P90 interact tightly with each other, forcing the release of SOs and occluding the binding site. At this stage the C-terminal domain of P1 has completed a hinge rotation of about 175° with respect to the “open” conformation, allowing the epitope of antibody P1/MCA4 to be fully exposed. Hinge movements of the C-terminal domain must be associated with displacements or distortions of the transmembrane helix, contiguous along the P1 sequence, which provides a way to communicate back and forth structural information between extra- and intra-cellular regions.

Binding of antibody P1/MCA4 to the C-terminal domain of P1 traps the Nap complex in the “closed” conformation, which explains why only moving cells are affected by the antibody (left images in **Figure 3)**. The increasing number of complexes in the closed, non-adherent conformation slows down the speed of gliding and weakens cell attachment.

Stage IV: Recovering the “open” conformation of the Nap complex. The “open” and “closed” conformations of the unbound adhesion complexes as described here so far, have the same molecular components. Assuming no other elements are at play, the least stable conformation must revert spontaneously towards the most stable, while the transition from the most towards the least stable will require energy. The “open” conformation, the most abundant in an *in situ* cryo-ET analysis (Aparicio et al., 2020), is likely the most stable, suggesting that the energy input is required for the “open” to “closed” transition associated with the power stroke and the straining of the internal structures (from Stage I to Stage III in **Figure 3**) ^11^. The force is likely generated by ATP hydrolysis at the back end of the TO ^27^ and transmitted to the adhesins through the contraction and extension of the TO internal rod structure that, by pulling the C-terminal domain of the P1 subunits, causes the transition between the conformations of the Nap complex.

Results provide now a consistent structural framework for the functioning of the Nap complex during the attachment/detachment cycle in which subunit P1, mainly its C-terminal domain, experiences extensive rearrangements pivoting around static P40/P90 subunits anchored by interactions of their Engelman motifs. Results also lay out a clear structural explanation for the efficient neutralization mechanism of antibodies such as P1/MCA4, binding to the dynamic and highly conserved C-terminal domain of P1, suggesting new approaches to confront human pathogens like *M. pneumoniae* and *M. genitalium*.

## Experimental Procedures

### Bacterial strains and tissue culture cells

*M. pneumoniae* M129 strain was grown in cell culture flasks containing SP4 or PPLO media and incubated at 37 °C with or without 5% CO_2_. Surface-attached mycoplasmas were harvested using a cell scraper and resuspended in medium or suitable buffers for experiments. When growing mycoplasma cells on IBIDI 8-well chamber slides, each well was seeded with about 10^5^ CFUs and incubated for 12-24 h in 200 μL SP4 supplemented with 3% gelatin. All *M. genitalium* strains were grown in SP-4 broth at 37°C in a 5% CO_2_ atmosphere in tissue culture flasks. SP-4 plates were prepared supplementing the medium with 0.8% agar (BD). Chloramphenicol (17 μg mL^−1^), puromycin (3 μg mL^−1^) or gentamycin (100 μg mL^−1^) were added for mutant selection.

*Escherichia coli* strain XL-1 Blue (Agilent, Santa Clara, USA) was used for cloning and plasmid propagation purposes. The strain was grown in Luria Bertani (LB) or LB agar plates containing 100 μg mL^−1^ ampicillin, 40 μg mL^−1^ X-Gal, and 24 μg mL^−1^ Isopropyl β-D-1-thiogalactopyranoside (IPTG) when needed.

NSI myeloma cells ^28^ were grown in RPMI 1640 medium supplemented with 10% FBS and 50 μg mL^-1^ gentamycin (complete RPMI). Hybridomas were selected in complete RPMI supplemented with HAT media and BM-Condimed (Sigma Aldrich).

### Cloning, expression and purification of P1 and P1 fragments

The coding sequences for constructs of P1 and the C-terminal domain of P1 were amplified from the synthetic clone of MPN_141 (P1) gene from *M. pneumoniae* (Genscript), using primers P1F and P1R for P1 (29-1521), P1Ct1400_F and P1R for P1 C-terminal (1400-1521), P1Ct1376_F and P1R for P1 C-terminal (1376-1521) (**Supplementary Table V**). The coding sequences for constructs of P40/P90 were amplified from the synthetic clone of MPN_142 (P40/P90) gene from *M. pneumoniae* (Genscript), using primers P40P90_F and P40P90_R (23-1114). The PCR fragments were cloned into the expression vector pOPINE (-) to add a C-terminal His-tag to the resulting constructs. Recombinant proteins were obtained after expression with B834 (DE3) cells (Merck), with IPTG at 0.8 mM and 22°C overnight. The cell pellets were lysed in 1xPBS at pH 7.4 and 40 mM Imidazole (binding buffer) and centrifuged at 20000 RPM at 4°C. Supernatant was charged into a HisTrap 5ml column (GE Healthcare) and eluted with a buffer containing 400 mM imidazole. Soluble aliquots were concentrated and loaded in a Superdex 200 GL 10/300 column (GE Healthcare) with a Tris·HCl 20 mM buffer (pH 7.4) and 150 mM NaCl.

### Production and validation of polyclonal antibodies

Polyclonal antisera anti-P1, anti-P1 fragments and anti-P90/P40 were prepared by immunizing BALB/C mice with the respective recombinant proteins and polypeptides described above. Sera were obtained by cardiac puncture of properly euthanized mice just before splenectomy and tittered using serial dilutions of each antigen. Titers of the different polyclonal sera were determined as the IC_50_ value from four parameter logistic plots and found to be approximately between 1/2500 and 1/4000.

### Production and sequencing of the monoclonal antibody P1/MCA4

A *p1* gene fragment was amplified by PCR from genomic DNA of *M. pneumoniae* M129 strain using primers P1F_2 and P1R_2 (**Supplementary Table V and VI)**. The amplified fragment was inserted into *Nco*I and *Xho*I site of pET-30c(+) expression vector, resulting the pP1-8 plasmid. *E. coli* BL21(DE3) was transformed by the pP1-8 for expression of a recombinant P1 peptide containing Ala1160 to Gln1518 of the P1 of M129 strain. The recombinant P1 was purified by His-tag affinity chromatography and used to immunize mice for production of anti-P1 monoclonal antibody. The monoclonal antibody P1/MCA4 was selected by ELISA screening and showed high specificity to P1 in Western blotting analysis and immunofluorescence microscopy of living *M. pneumoniae* cells. The amino acid sequence of P1/MCA4 was determined at GenScript (Tokyo, Japan) by cloning and sequencing of the cDNAs of antibody mRNA from P1/MCA4 producing hybridoma cell. The obtained sequences for a single H chain and two L chain of the P1/MCA4 (**Supplementary** Figure 1) were deposited into the DDBJ/ENA/GenBank databases under the accession numbers LC600310, LC600311 and LC600312.

### Purification of the Fab fragment

Monoclonal antibody P1/MCA4 against P1 was diluted up to 2 mg mL^-1^ in 1xPBS buffered at pH 8.0 and mixed with 0.1mg/ml of Papain (previously activated in 20mM L-cysteine, 20 mM EDTA and 1xPBS pH: 7.0) in a relation 1:40 (Protein:Papain) and incubated during 4 hours at 37°C. To stop digestion, 10% of the final volume of Iodine acetamide was added. The digested solution was loaded onto HiTrap Protein G 5ml with Binding buffer (20 Mm Na_2_HPO_4_ pH:7.0) and Elution buffer (0.1 M Glycine·HCl pH 2.7). Soluble aliquots were concentrated and loaded in a Superdex 75 GL 10/300 column (GE Healthcare) with a Tris·HCl 20 mM pH 7.4 and 150 mM NaCl buffer. Complexes of P1-mAb and P1-Fab were formed by mixing both proteins in a ratio 2:1 and 1:1, respectively.

### Epitope mapping of Monoclonal antibody P1/MCA4

For epitope mapping of P1/MCA4, a series of plasmids that express recombinant P1 peptides were constructed (**Supplementary Table VI**). These plasmids were generated by partial deletion of the pP1-8 plasmid using specific primer sets and PrimeSTAR Mutagenesis Basal kit (Takara Bio, Shiga, Japan). Recombinant P1 peptides were produced in *E. coli* BL21(DE3) harboring the plasmids, separated by 10-20% gradient-gel SDS-PAGE, and transferred to nitrocellulose membranes. Recombinant P1 peptides on membranes were reacted with 0.5 µg/ml P1/MCA4. After wash of the membranes, binding of the P1/MCA4 to P1 peptide was detected by anti-mouse IgG, HRP conjugate secondary antibody (Promega, Madison, WI, USA) and EzWestBlue W substrate (Atto, Tokyo, Japan).

### Size Exclusion Chromatography and Multi Angle Light Scattering (SEC-MALS)

Molecular weights were measured using a Superdex 200 10/300 GL (GE HEalthcare) column in a Prominence liquid chromatography system (Shimadzu) connected to a DAWN HELEOS II multi-angle light scattering (MALS) detector and an Optilab T-REX refractive index (dRI) detector (Wyatt Technology). ASTRA 7 software (Wyatt Technology) was used for data processing and result analysis. A dn/dc value of 0.185 mL g^-1^ (typical of proteins) was assumed for calculations.

### Single-particle cryo-EM of the binary complex

To prevent preferred orientations, Octylthioglucoside (OG) was added to solution of purified P1-Fab (P1/MCA4) complex to adjust the final concentration of OG to 0.9% (w/l). 2.6 μL of sample solution was applied to a glow-discharged Au-coated Quantifoil holey carbon grid (R1.2/1.3, Cu, 200 mesh), blotted for 4 sec at 4 °C in 100% humidity and plunged into frozen liquid ethane using a Vitrobot Mark IV (Thermo Fisher Scientific). The grid was inserted into a CRYO ARM 300 (JEOL) operating at an acceleration voltage of 300 kV. CryoEM images were recorded with a K3 direct electron detector (Gatan) in CDS mode with an energy filter at a slit width of 20 eV. Data were automatically collected using the SerialEM software (https://bio3d.colorado.edu/SerialEM/) at a physical pixel size of 0.49 Å with 50 frames at a dose of 2.0 e^−^/ Å ^2^ per frame and an exposure time of 2.86 sec per movie with a defocus ranging from -0.5 to -1.5 μm. A total of 27,122 movies were collected for P1-Fab complex.

The movie frames were subjected to beam-induced movement correction using MotionCor2.1 ^29^ and contrast transfer function (CTF) was evaluated using Gctf ^30^. Approximately 2,000 particles were manually selected from 20 micrographs to perform two-dimensional (2D) classification. Using a good 2D class average image, a total of 4,312,408 particle images were automatically picked and 2D classifications were performed using RELION-3.1 ^31^. A total of 1,252,491 particles were selected for building the initial model of the P1-Fab complex using cryoSPARC2 ^32^ and subjected to 3D classification into 8 classes using RELION-3.1. The selected particles were re-extracted at a pixel size of 0.49 Å and subjected to three 3D refinement, two CTF refinement and Bayesian polishing. After selection of particles using no-alignment 3D classification, a total of 352,039 particles were subjected to two 3D refinement and CTF refinement. The final 3D refinement and post-processing yielded maps with global resolutions of 2.39 Å, according to the 0.143 criterion of the Fourier shell correlation **(Supplementary Table II)**. Local resolution was estimated using RELION-3.1. Processing strategy is described in **Supplementary** Figure 4.

### Model building and refinement of the binary complex

The model of the P1-Fab complex was built based on the cryoEM density map. Previous published crystal structure of P1 (PDB ID: 6RC9) was docked into the EM density map using UCSF Chimera ^32^. N-domain, C-domain of P1 and Fab were manually repositioned and refined iteratively using COOT ^33^ and PHENIX real space refinement ^34^. The statistics of the 3D reconstruction and model refinement are summarized in **Supplementary Table II**. The final refined structure has been deposited in the PDB with code 8ROR.

### Single-particle cryo-EM of the ternary complex

2.6 μL of sample solution was applied to a glow-discharged Quantifoil holey carbon grid (R1.2/1.3, Cu, 200 mesh), blotted for 4 sec at 4°C in 100% humidity and plunged into frozen liquid ethane using a Vitrobot Mark IV. The grid was inserted into a CRYO ARM 300 operating at an acceleration voltage of 300 kV. CryoEM images were recorded with a K3 direct electron detector in CDS mode with an energy filter at a slit width of 20 eV. Data were automatically collected using the SerialEM software (https://bio3d.colorado.edu/SerialEM/) at a physical pixel size of 0.87 Å with 40 frames at a dose of 2.0 e^−^/ Å ^2^ per frame and an exposure time of 3.3 sec per movie with a defocus ranging from -0.8 to -1.8 μm. A total of 7,938 movies were collected for P40/P90-P1-Fab complex.

The movie frames were subjected to beam-induced movement correction using MotionCor2.1 and CTF was evaluated using Gctf. Using blob picker program of cryoSPARC2, a total of 9,565,198 particle images were automatically picked and 2D classifications were performed using cryoSPARC2. A total of 1,070,120 particles were selected for building the initial model of the P40/P90-P1-Fab complex using cryoSPARC2. However, due to preferred orientation, an accurate initial structure could not be obtained. Therefore, we did not perform 3D reconstruction for high-resolution structure determination, but performed 3D refinement to confirm how P40/P90 and Fab are bound to P1. The selected particles were re-extracted at a pixel size of 0.87 Å and subjected to 3D refinement. The final 3D refinement and post-processing yielded maps with global resolutions of below 10 Å, according to the 0.143 criterion of the Fourier shell correlation **(Supplementary Table II)**.

### DNA manipulation and primers

Plasmid DNA was purified using GeneJET Plasmid Miniprep Kit (Thermo Fisher Scientific). PCR products and DNA fragments were recovered from agarose gels using NucleoSpin Gel and PCR Clean-up Kit (Macherey-Nagel, Düren, Germany), and digested using the corresponding restriction enzymes (Thermo Fisher Scientific) when necessary. For transformation of *M. genitalium*, plasmids were purified using the GenElute HP Midiprep Kit (Sigma-Aldrich, St. Louis, USA) following the manufacturer’s instructions. All primers used in this study are listed in **Supplementary Table VII**.

### M. genitalium mutant strains

The suicide plasmid pBEΔMG_191/MG_192 was designed to generate a G37 *M. genitalium* MG_191 and MG_192 null mutant strain by gene replacement. First, a 1 kb flanking upstream region (UR) to the MG_191 gene was PCR-amplified using the MgParUp-F and MgParUp-R primers. Similarly, a 1 kb flanking downstream region to the MG_192 gene was PCR-amplified with the MgParDw-F and MgParDw-R primers. In parallel, the a lox version of the CmM438 selectable marker ^35^ was PCR-amplified with the Lox71p438-Fwd-XhoI and CatLox66-Rev-BamHI primers, obtaining the CmM438 flanked with the lox61 and lox71 sequences. Then, the UR and DR PCR products were joined by PCR with the MgParUp-F and MgParDw-R primers. The resulting PCR product was cloned into an EcoRV-digested pBE plasmid ^36^. Finally, the resulting CmM438 amplicon was digested with XhoI and BamHI restriction enzymes and ligated into the similarly digested pBE containing the UR and DR regions obtained before.

This plasmid was used to obtain the G37ΔAdh chloramphenicol resistant mutant strain. Electroporation of this mutant with the pCre plasmid ^37^ that contains the Cre recombinase allowed us to obtain the G37ΔAdh strain, free of any antibiotic selectable marker. This strain was used as the recipient strain to transform all the pMTnPac plasmids generated in this study.

MG_191 and MG_192 genes from the chromosome of *M. genitalium* G37 strain were amplified by PCR using COMmg191-F-ApaI and COMmg191/192-R-SalI primers. The resulting PCR product was cloned into an EcoRV-digested pBE plasmid to create the pBE-COMP140P110. At the same time, the amplicon was digested with the ApaI and SalI restriction enzymes and ligated to a similarly digested pMTnPac plasmid to create the pTnPacCOMP140P110. This plasmid was used to reintroduce the wild-type alleles of the MG_191 and MG_192 genes to the G37ΔAdh mutant. The COMmg192-F (ApaI) primer includes the upstream region (70 nucleotides) of the MG_191 gene, which contains a strong promoter identified in a previous study ^38^. This promoter was used to drive the transcription of the transposon-encoded copy of the MG_191 and MG_192 genes in all mutants.

To generate P140 and P110 variants carrying specific mutations in the Engelman motifs, the target mutations in the MG_191 and or MG_192 genes were introduced by Exsite-PCR using the pBE-COMP140P110 as a template and the specific primers for each mutant (**Supplementary Table VII)**. Then, plasmids containing the P140 and P110 alleles with the desired mutations were re-ligated and mutant P140 and P110 alleles excised by digestion with ApaI and SalI restriction enzymes. Finally, DNA fragments with mutant alleles were ligated into a pMTnPac plasmid previously digested with ApaI and SalI, to generate the corresponding pMTnPacCOMP140P110 plasmid series.

Sequencing analysis of the different TnPacCOMP140P110 constructs using primers Tnp3, RTPCR192-F, RTPCR192-R, and PacUp, ruled out the presence of additional mutations in the MG_191 and MG_192 sequences. These plasmids were transformed into the *M. genitalium* G37ΔAdh null mutant to create the different P140 and P110 variant strains. Identification of the minitransposon insertion site in the individual clones was done by sequencing using the PacDown primer and chromosomal DNA as a template.

### Transformation and screening

*M. genitalium* G37ΔAdh null mutants were transformed by electroporation using 10 μg of plasmid DNA of the different minitransposons or 30 μg when using plasmids for gene replacement experiments, as previously described ^36,39^. Puromycin-resistant colonies were picked, propagated, and stored at –80 °C. For screening purposes, strains were further propagated in antibiotic supplemented SP4 medium in 25 cm^2^ tissue culture flasks. To obtain crude lysates from grown cultures, SP4 medium was removed and cells were lysed using 50 μL of Lysis Buffer (0.1 M Tris-HCl pH 8.5, 0.05% Tween-20, and 250 μg mL^-1^ Proteinase K) and incubated for 1 h at 37 °C. Then, Proteinase K was inactivated at 96 °C for 10 min. *M. genitalium* lysates were screened by PCR or direct Sanger sequencing using the primers in **Supplementary Table VII**.

### Sequencing reactions

DNA sequencing reactions were performed using BigDye® v3.1 Cycle Sequencing kit using 2.5 μL of genomic DNA or *M. genitalium* lysate, following manufacturer’s instructions (Thermo Fisher Scientific). All reactions were analysed in an ABI PRISM 3130xl Genetic Analyzer at the Servei de Genòmica i Bioinformàtica (UAB).

### SDS-PAGE and Western Blotting

Whole-cell lysates were obtained from mid-log phase cultures grown in 75 cm^2^ flasks. Protein concentration was determined with the Pierce^TM^ BCA Protein Assay Kit (Thermo Fisher Scientific), and similar amounts of total protein were separated by SDS-PAGE following standard procedures.

For specific detection of *M. genitalium* proteins, SDS-PAGE gels were electrotransferred to PVDF membranes by a Semi-dry transfer System, using cold Towbin buffer (25 mM Tris, 192 mM Glycine pH 8.3-8.4, 20% Methanol (v/v)) following the standard protocols ^40,41^. Membranes were probed with the appropriate primary antibody in blocking solution, which was then detected using a secondary antibody conjugated with horseradish peroxidase (Bio-Rad). Antibodies used in this study are listed in **Supplementary Table VIII.** Bioluminescence reaction was catalyzed with *Luminata Forte^TM^ Western HRP substrate* (Merk Millipore). Visualization and image optimization was performed in a VersaDoc Imaging System (Bio-Rad) using QuantityOne^®^ software (Bio-Rad).

### Quantitative hemadsorption assay

Hemadsorption was quantified using flow cytometry as previously described ^42^ with few modifications. We used 10^9^ mycoplasma cells for the hemadsorption assay. Fluorescence-activated cell sorting (FACS) data were acquired using a FACSCalibur (Becton Dickinson, Franklin Lakes, USA) equipped with an air-cooled 488 nm argon laser and a 633 nm red diode laser and analysed with the CellQuest-Pro and FACSDiva software (Becton Dickinson). Binding of mycoplasma cells to red blood cells was modelled in an inverse Langmuir isothermal kinetic function:

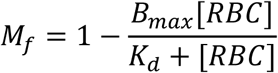

Plots represent the best-fitting curves to a series of hemadsorption measurements obtained from at least two biological repeats for each strain. We performed a double-gating strategy, using a preliminary FL3-H/FL2-H gate following an SSC-H/FL1-H gate.

### Time lapse micro-cinematography

Effects of antibody sera on mycoplasma cells were investigated by time lapse cinematography of *M. pneumoniae* cells growing in IBIDI 8-well chamber slides using SP4 medium supplemented with 3. Prior to observation, the medium was replaced with fresh SP4 supplemented with 3% gelatin and pre-warmed at 37°C. After incubation for 10 minutes at 37 °C and 5% CO_2_, the slide was placed in a Nikon Eclipse TE 2000-E inverted microscope equipped with a Microscope Cage Incubation System (Okolab) at 37 °C. Images were captured at 0.5 s intervals for a total observation time of 10 min, and the different antibodies were dispensed directly into the wells after the first 50 seconds of observation. Frequencies of motile cells and detached cells before adding the different antibodies were calculated from the images collected between 0 and 50 seconds of observation. Frequencies of motile cells and detached cells after adding the different antibodies were calculated from the pictures collected between 50 seconds and 10 minutes of observation.

### Immunofluorescence microscopy

The immunofluorescence staining of mycoplasma cells on chamber slides was similar to previously described ^9,20^, with several modifications. All mycoplasma cells were washed with PBS containing 0.02% Tween 20 (PBS-T) except cells from *M. genitalium* G37ΔAdh strain and Engelman motif mutant strains, which were washed in PBS containing 136 mM sucrose (PBS-Suc), prewarmed at 37 °C. Then, each well was fixed with 200 μL of 3% paraformaldehyde (wt/vol) and 0.1% glutaraldehyde. Cells were washed three times with PBS-T (or PBS-Suc), and slides were immediately treated with 3% BSA in PBS-T (blocking solution) for 30 minutes. The blocking solution was removed and each well was incubated for 1 hour with 200 μL of the primary antibodies diluted in blocking solution. We have used a 1/2000 dilution for all polyclonal sera. Wells were washed three times with PBS-T or PBS-Suc and incubated for 1 hour with a 1/2000 dilution of a goat anti-mouse Alexa 555 secondary antibody (Invitrogen) in blocking solution. Wells were then washed three times with PBS-T or PBS-Suc and incubated for 20 minutes with 100 μL of a solution of Hoechst 33342 10 μg μL^-1^ in PBS-T or PBS-Suc. Wells were finally washed once with PBS-T and replenished with 200 μL of PBS-T. Cells were examined by phase contrast and epifluorescence in an Eclipse TE 2000-E inverted microscope (Nikon). Phase contrast images, 4’,6-diamidino-2-phenylindole (DAPI, excitation 387/11 nm, emission 447/60 nm) and Texas Red (excitation 560/20 nm, emission 593/40 nm) epifluorescence images were captured with an Orca Fusion camera (Hamamatsu) controlled by NIS-Elements BR software (Nikon).

### Scanning Electron Microscopy

For the Scanning Electron Microscopy (SEM) analysis, the *M. genitalium* cultures were grown to mid-long phase over glass and round coverslips in a 24-well plate with fresh SP4. For non-adherent mutant strains, the coverslips were treated in advance with 0.2 mg/mL Poly-L-Lysine to allow the cells to stick to the surface of the glass. The inoculum depended on the stock concentration and usually ranged between 5-10 µL of *M. genitalium* strains. The *Mge*-WT and the mutant strains were grown overnight and washed twice with 1 mL 0.1M PB. Then, the cells were fixed with 2% paraformaldehyde and 2.5% glutaraldehyde (v/v) dissolved in 0.1M PB for two hours in the dark and room temperature. Samples were then washed again with 0.1M PB and dehydrated sequentially with increasing ethanol solutions (25, 50, 75 and 100% ethanol), 10 minutes for each solution. Once in 100% ethanol, samples were sent to Servei de Microscòpia (UAB) and immediately critical-point dried in a *K850 Critical Point Dryer* (Emitech) and then they were sputter coated with gold. Samples were observed using a MERLIN FE-SEM microscope (ZEISS).

## Supporting information

Video III

## Acknowledgements

We are most thankful to Dr. X. Daura for his help in the epitope prediction study. We also thank L. Company and I. Fernández-Vidal for their support during MALS measurements, to A. Iborra (Servei de Cultius Cellulars, Anticossos Citometria, UAB) for his assistance with immunizing mice, and D. Nakane (The University of Electro-Communications) for assistance of microscopic observation of *M. pneumoniae*. J.R.T. and D.V. were supported by an EMBO (under grant agreement ALTF 145-2021) and a Margarita Salas (Barcelona University) postdoctoral fellowships, respectively. E.M. was supported by grants RYC2018-025295-I and PID2020-120098GA-I00 and I. F.-J. P. by grants PID2021-125632OB-C21 and -C22 from the Spanish Ministry of Science and Innovation. This work was also supported by a Grants-in-Aid for Scientific Research (15H01337 and 22K07063) from the Ministry of Education, Culture, Sports, Science, and Technology of Japan to T.K.

