## Supplementary figures for "Dynamics of the adhesion complex of the human pathogens *Mycoplasma pneumoniae and Mycoplasma genitalium*"

### P1/MCA4 H chain

DDBJ/ENA/GenBank accession no. LC600310

Signal peptide -FR1-CDR1-FR2-CDR2-FR3-CDR3-FR4-constant region

```
atgggatggagctggatctttctcttcctcctgtcaggaactgcaggtgtccactctgaggtccagctgcaacagtctggacctgaactggtgaagcctggaacttcaatgaagatatcc   120
M G W S W I F L F L L S G T A G V H S E V Q L Q Q S G P E L V K P G T S M K I S   40
tgcaaggcttctggttactcattcactggctacaccatgaactgggtgaagcagagccatggaaagagccttgagtggattggacttattaatccttacaatggtggtactaactacaac   240
C K A S G Y S F T G Y T M N W V K Q S H G K S L E W I G L I N P Y N G G T N Y N   80
cagaagttcaggggcacggccacattaactgtagacaagtcatccagcacagcctacatggagctcctcagtctgacatctgaggactctgcagtctattactgtgcaaggtcgaactat   360
Q K F R G T A T L T V D K S S S T A Y M E L L S L T S E D S A V Y Y C A R S N Y   120
gcttacgacctattgatggactactggggtcaaggaacctcagtcaccgtctcctcagccaaaacgacacccccatctgtctatccactggcccctggatctgctgcccaaactaactcc   480
A Y D L L M D Y W G Q G T S V T V S S A K T T P P S V Y P L A P G S A A Q T N S   160
atggtgaccctgggatgcctggtcaagggctatttccctgagccagtgacagtgacctggaactctggatccctgtccagcggtgtgcacaccttcccagctgtcctgcagtctgacctc   600
M V T L G C L V K G Y F P E P V T V T W N S G S L S S G V H T F P A V L Q S D L   200
tacactctgagcagctcagtgactgtcccctccagcacctggcccagcgagaccgtcacctgcaacgttgcccacccggccagcagcaccaaggtggacaagaaaattgtgcccagggat   720
Y T L S S S V T V P S S T W P S E T V T C N V A H P A S S T K V D K K I V P R D   240
tgtggttgtaagccttgcatatgtacagtcccagaagtatcatctgtcttcatcttccccccaaagcccaaggatgtgctcaccattactctgactcctaaggtcacgtgtgttgtggta   840
C G C K P C I C T V P E V S S V F I F P P K P K D V L T I T L T P K V T C V V V   280
gacatcagcaaggatgatcccgaggtccagttcagctggtttgtagatgatgtggaggtgcacacagctcagacgcaaccccgggaggagcagttcaacagcactttccgctcagtcagt   960
D I S K D D P E V Q F S W F V D D V E V H T A Q T Q P R E E Q F N S T F R S V S   320
gaacttcccatcatgcaccaggactggctcaatggcaaggagttcaaatgcagggtcaacagtgcagctttccctgcccccatcgagaaaaccatctccaaaaccaaaggcagaccgaag   1080
E L P I M H Q D W L N G K E F K C R V N S A A F P A P I E K T I S K T K G R P K   360
gctccacaggtgtacaccattccacctcccaaggagcagatggccaaggataaagtcagtctgacctgcatgataacagacttcttccctgaagacattactgtggagtggcagtggaat   1200
A P Q V Y T I P P P K E Q M A K D K V S L T C M I T D F F P E D I T V E W Q W N   400
gggcagccagcggagaactacaagaacactcagcccatcatggacacagatggctcttacttcgtctacagcaagctcaatgtgcagaagagcaactgggaggcaggaaatactttcacc   1320
G Q P A E N Y K N T Q P I M D T D G S Y F V Y S K L N V Q K S N W E A G N T F T   440
tgctctgtgttacatgagggcctgcacaaccaccatactgagaagagcctctcccactctcctggtaaatga   1392
C S V L H E G L H N H H T E K S L S H S P G K *   463
```

### P1/MCA4 L chain 1 (L1)

DDBJ/ENA/GenBank accession no. LC600311

Signal peptide -FR1-CDR1-FR2-CDR2-FR3-CDR3-FR4-constant region

```
atgaagttgcctgttaggctgttggtgctgatgttctggattcctgcttccagcagtgatgttttgatgacccaaactccactctccctgcctgtcagtcttggagatcaagcctccatc   120
M K L P V R L L V L M F W I P A S S S D V L M T Q T P L S L P V S L G D Q A S I   40
tcttgcagatttagtcagaccattgtacatagtaatggagccacctatttagaatggtacctgcagagaccaggccagtctccaaagctcctgatctacaaagtttccaaccgattttct   240
S C R F S Q T I V H S N G A T Y L E W Y L Q R P G Q S P K L L I Y K V S N R F S   80
ggggtcccagacaggttcagtggcagtggatcagggacagatttcacactcaaaatcagcagagtggaggctgaggatctgggagtttattactgctttcaaggttcacatgttccgtgg   360
G V P D R F S G S G S G T D F T L K I S R V E A E D L G V Y Y C F Q G S H V P W   120
acgttcggtggaggcaccaagctggaaatcaaacgggctgatgctgcaccaactgtatccatcttcccaccatccagtgagcagttaacatctggaggtgcctcagtcgtgtgcttcttg   480
T F G G G T K L E I K R A D A A P T V S I F P P S S E Q L T S G G A S V V C F L   160
aacaacttctaccccaaagacatcaatgtcaagtggaagattgatggcagtgaacgacaaaatggcgtcctgaacagttggactgatcaggacagcaaagacagcacctacagcatgagc   600
N N F Y P K D I N V K W K I D G S E R Q N G V L N S W T D Q D S K D S T Y S M S   200
agcaccctcacgttgaccaaggacgagtatgaacgacataacagctatacctgtgaggccactcacaagacatcaacttcacccattgtcaagagcttcaacaggaatgagtgttag   717
S T L T L T K D E Y E R H N S Y T C E A T H K T S T S P I V K S F N R N E C *   238
```

### P1/MCA4 L chain 2 (L2)

DDBJ/ENA/GenBank accession no. LC600312

Signal peptide -FR1-CDR1-FR2-CDR2-FR3-CDR3-FR4-constant region

```
atgagtcctgcccagttcctgtttctcttagtgctctggattcgggaaaccaacggtgatgttgtgatgacccagactccactcactttgtcggttaccattggacaaccagcctccatc   120
M S P A Q F L F L L V L W I R E T N G D V V M T Q T P L T L S V T I G Q P A S I   40
tcttgcaagtcaagtcagagcctcttagatagtgatggaaagacatatttgaattgcttcttacagaggccaggccagtctccaaatcgcctgatctatctggtgtctagactggactct   240
S C K S S Q S L L D S D G K T Y L N C F L Q R P G Q S P N R L I Y L V S R L D S   80
ggagtccctgacaggttcactggcagtggatcagggacagatttcacactgaaaatcagcagagtggaggctgaggatttgggagtttattattgctgtcaagttacacagtggacgttc   360
G V P D R F T G S G S G T D F T L K I S R V E A E D L G V Y Y C C Q V T Q W T F   120
ggtggaggcaccaagctggaaatcaaacgggctgatgctgcaccaactgtatccatcttcccaccatccagtgagcagttaacatctggaggtgcctcagtcgtgtgcttcttgaacaac   480
G G G T K L E I K R A D A A P T V S I F P P S S E Q L T S G G A S V V C F L N N   160
ttctaccccaaagacatcaatgtcaagtggaagattgatggcagtgaacgacaaaatggcgtcctgaacagttggactgatcaggacagcaaagacagcacctacagcatgagcagcacc   600
F Y P K D I N V K W K I D G S E R Q N G V L N S W T D Q D S K D S T Y S M S S T   200
ctcacgttgaccaaggacgagtatgaacgacataacagctatacctgtgaggccactcacaagacatcaacttcacccattgtcaagagcttcaacaggaatgagtgttag   711
L T L T K D E Y E R H N S Y T C E A T H K T S T S P I V K S F N R N E C *   236
```

### Alignment of L1 and L2 amino acid sequences

```
L1  MKLPVRLLVLMFWIPASSSDVLMTQTPLSLPVSLGDQASISCRFSQTIVHSNGATYLEWYLQRPGQSPKLLIYKVSNRFSGVPDRFSGSGSGTDFTLKISRVEAEDLGVYYCFQGSHVPW  12 0
    |    |  |  ||     || |||||| | |  |  |||||  ||     | | |||   ||||||||  ||| ||   ||||||| ||||||||||||||||||||||||| |      |
L2  MSPAQFLFLLVLWIRETNGDVVMTQTPLTLSVTIGQPASISCKSSQSLLDSDGKTYLNCFLQRPGQSPNRLIYLVSRLDSGVPDRFTGSGSGTDFTLKISRVEAEDLGVYYCCQVTQ --W  118

    TFGGGTKLEIKRADAAPTVSIFPPSSEQLTSGGASVVCFLNNFYPKDINVKWKIDGSERQNGVLNSWTDQDSKDSTYSMSSTLTLTKDEYERHNSYTCEATHKTSTSPIVKSFNRNEC  238
    ||||||||||||||||||||||||||||||||||||||||||||||||||||||||||||||||||||||||||||||||||||||||||||||||||||||||||||||||||||||
    TFGGGTKLEIKRADAAPTVSIFPPSSEQLTSGGASVVCFLNNFYPKDINVKWKIDGSERQNGVLNSWTDQDSKDSTYSMSSTLTLTKDEYERHNSYTCEATHKTSTSPIVKSFNRNEC  236
```

**Supplementary Figure 1. Nucleotides and amino-acids sequences corresponding to the Heavy and the (two) Light chains from Mab P1/MCA4**

**a**

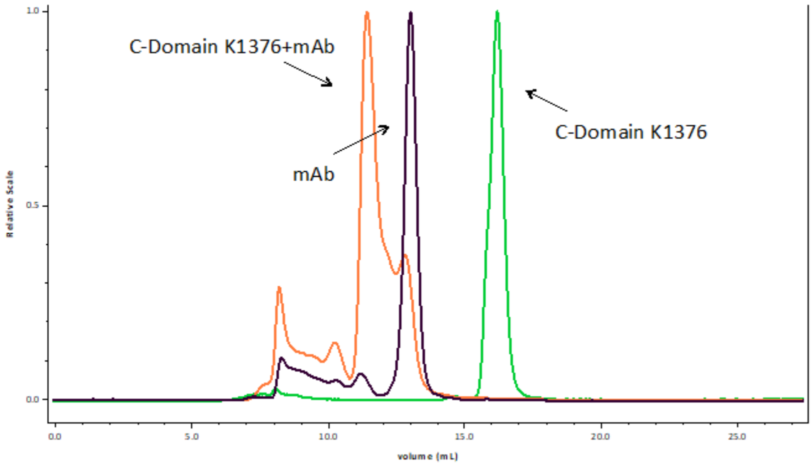


| Peak | Mw (KDa) | Mass Fraction (%) |
|---|---|---|
| C-Domain K1376 | 18.92 ± 0.07 | 100.00 |
| mAb | 148.44 ± 0.20 | 100.00 |
| C-Domain K1376+mAb | 189.53 ± 0.21 | 67.10 |

**b**

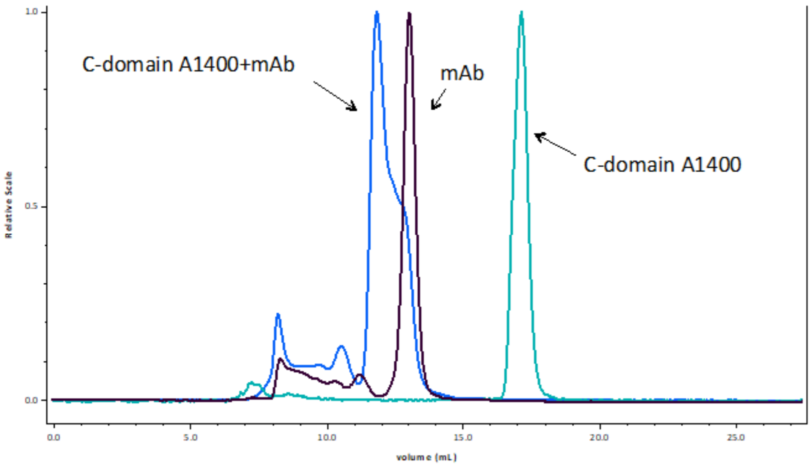


| Peak | Mw (KDa) | Mass Fraction (%) |
|---|---|---|
| C-Domain A1376 | 15.79 ± 0.09 | 100.00 |
| mAb | 148.44 ± 0.20 | 100.00 |
| C-Domain A1376+ mAb | 179.44 ± 0.21 | 57.07 |

**Supplementary Figure 2. Analysis by MALS of samples containing Mab P1/MCA4 and the C-terminal domain constructs from P1.** In each of the two samples containing the Mab P1/MCA4 and a construct of the C-terminal domain from P1 is clear the presence of a complex with molecular weight of ~179.4 and 189.5 kDa, for constructs A1400 (a) and K1376 (b), respectively**.**

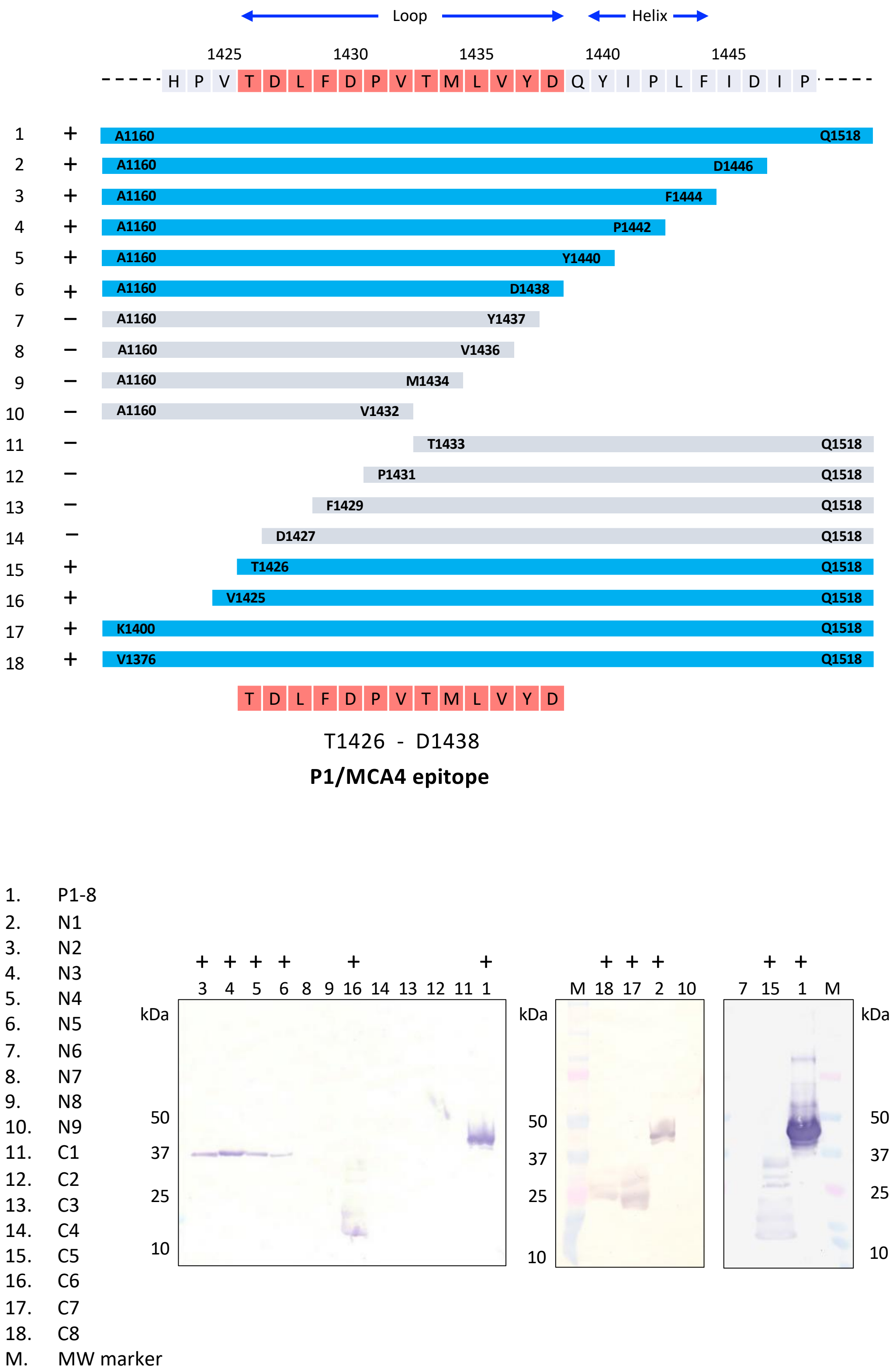


**Supplementary Figure 3. P1 epitope mapping of Mab P1/MCA4.**

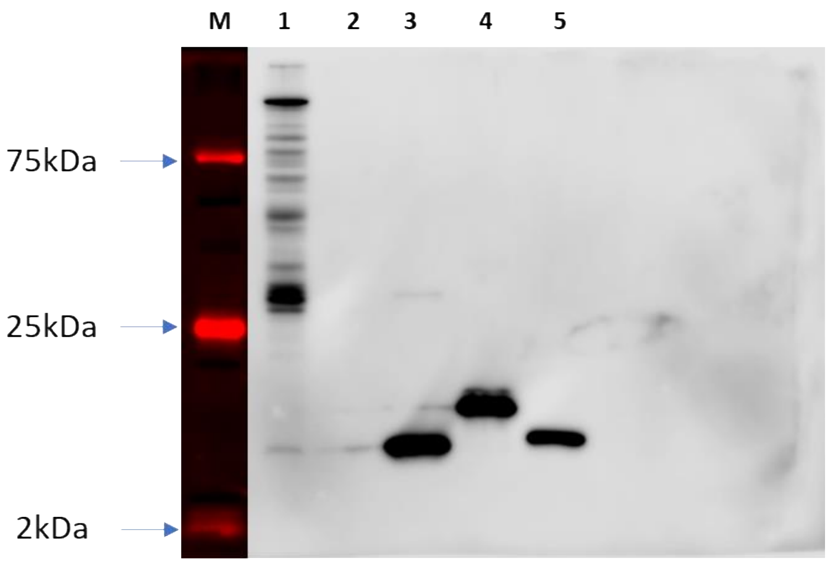


**Supplementary Figure 4. Western blotting analysis performed using Mab P1/MCA4 and different constructs from P1 (*M. pneumoniae*) and from P140 (*M. genitalium*).** 1) *Mpn*-WT (MPN129); 2) P1Glob. Thr29-Ala1375; 3) P1 C-terminal A1400-D1521; 4) P1 C-terminal K1376-D1521; 5) C-terminal P140 S1244-D1351. The key peptide in the epitope of P1 (1426 TDLFDPVTMLVYD 1438) presents a high sequence identity with the corresponding peptide in P140 (1270 TELFDPNTMFVYD 1282).

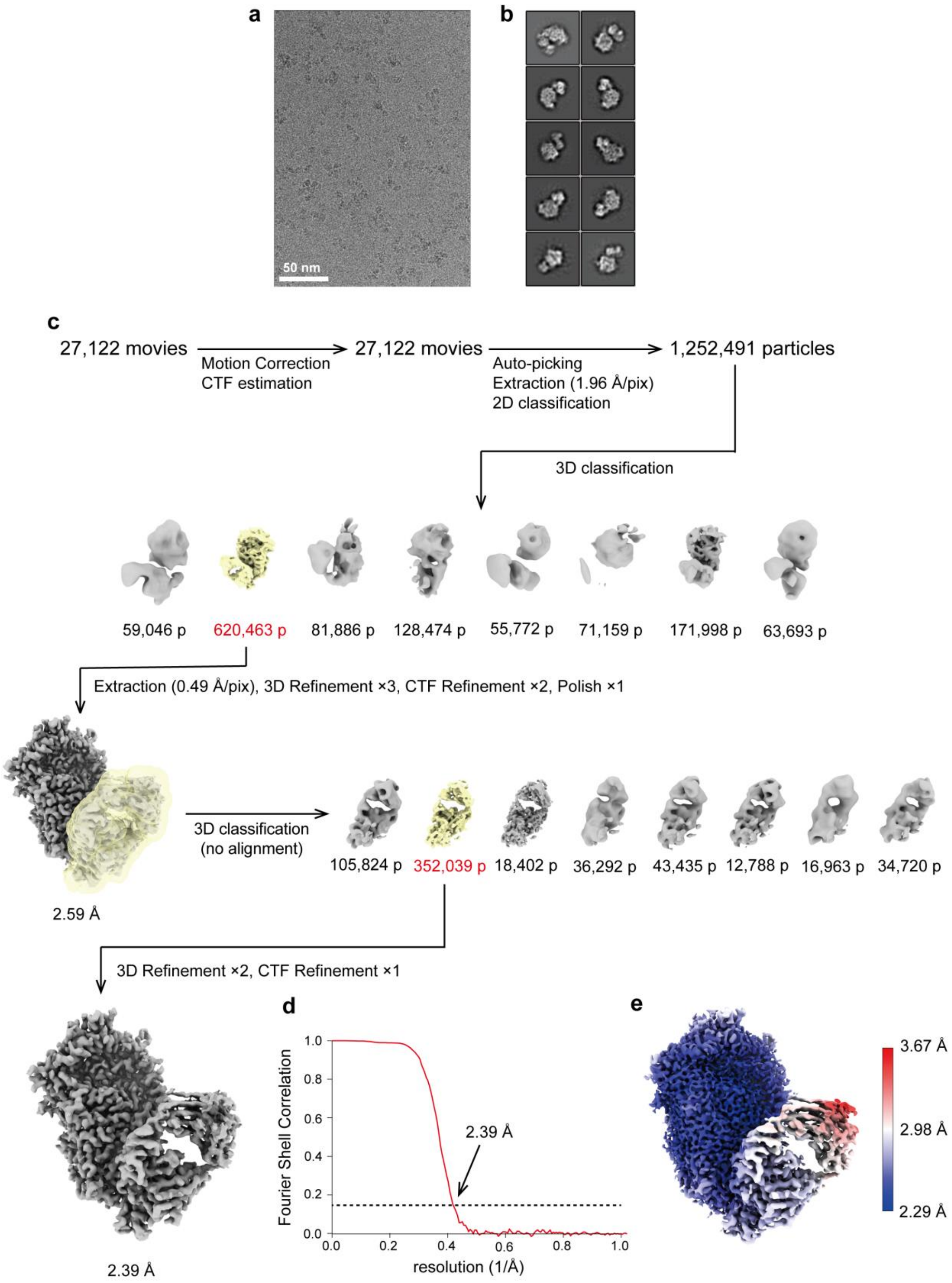


**Supplementary Figure 5**. **Cryo-EM structure determination of the P1-Fab(P1/MCA4) complex.**

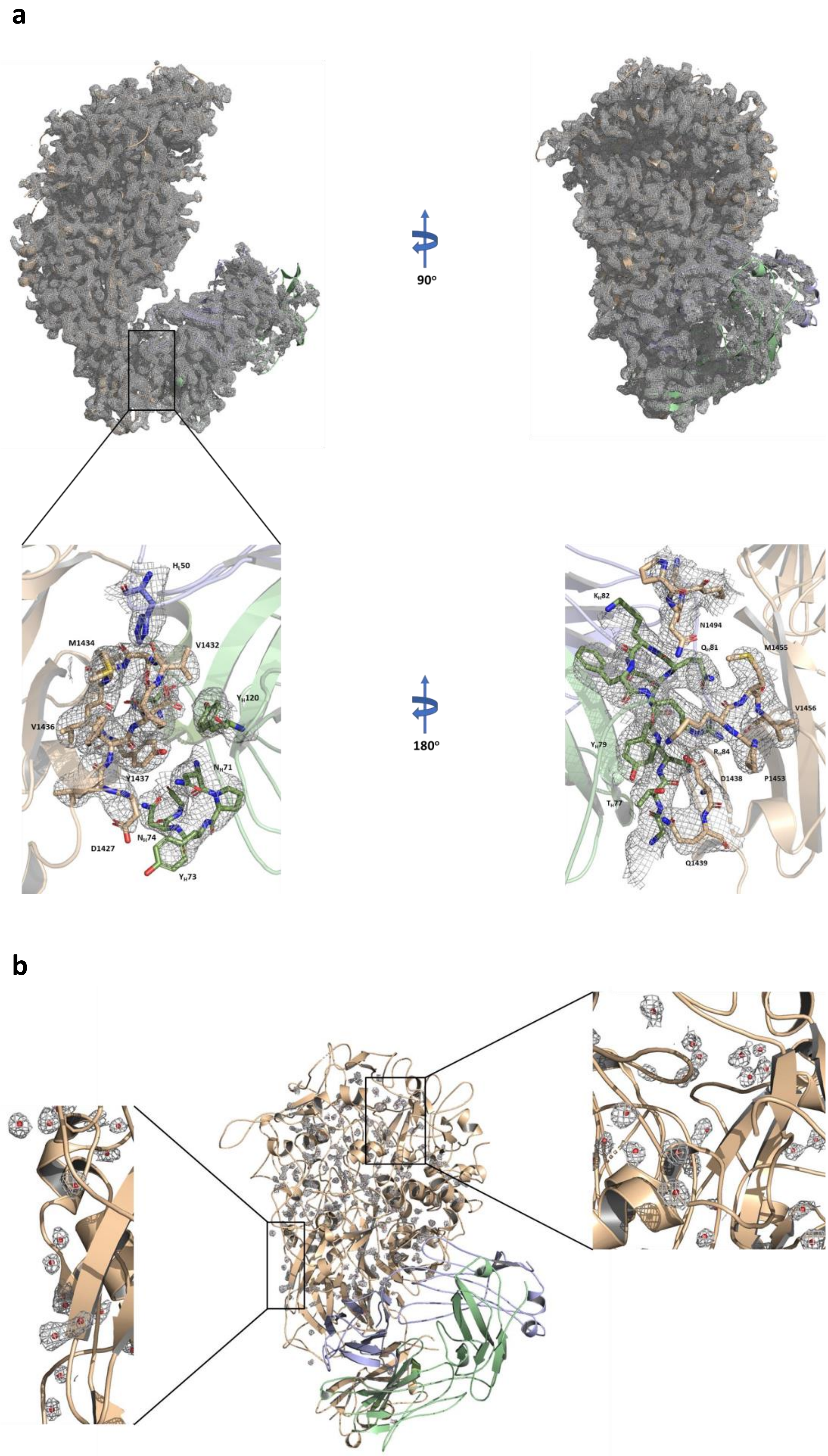
a
90°
$H_L$50
M1434
V1432
$Y_H$120
V1436
Y1437
$N_H$71
D1427
$N_H$74
$Y_H$73
180°
$K_H$82
N1494
$Q_H$81
M1455
V1456
$Y_H$79
$R_H$84
D1438
P1453
$T_H$77
Q1439
b

**Supplementary Figure 6. Map quality of the P1-Fab(P1/MCA4) complex: Identification of solvent molecules. a)** Two 90° apart views of the whole cryo-EM map of the P1-Fab(P1/MCA4) complex. The map is crispy, with well-defined side chains, for most of P1 and also for the Fab variable module ($V_L$-$V_H$). Inset shows the map at the epitope-paratope interface (same color code as in Figure 1a). **b)** The quality of the map allowed the identification of a large number of solvent molecules in the N-terminal domain of P1.

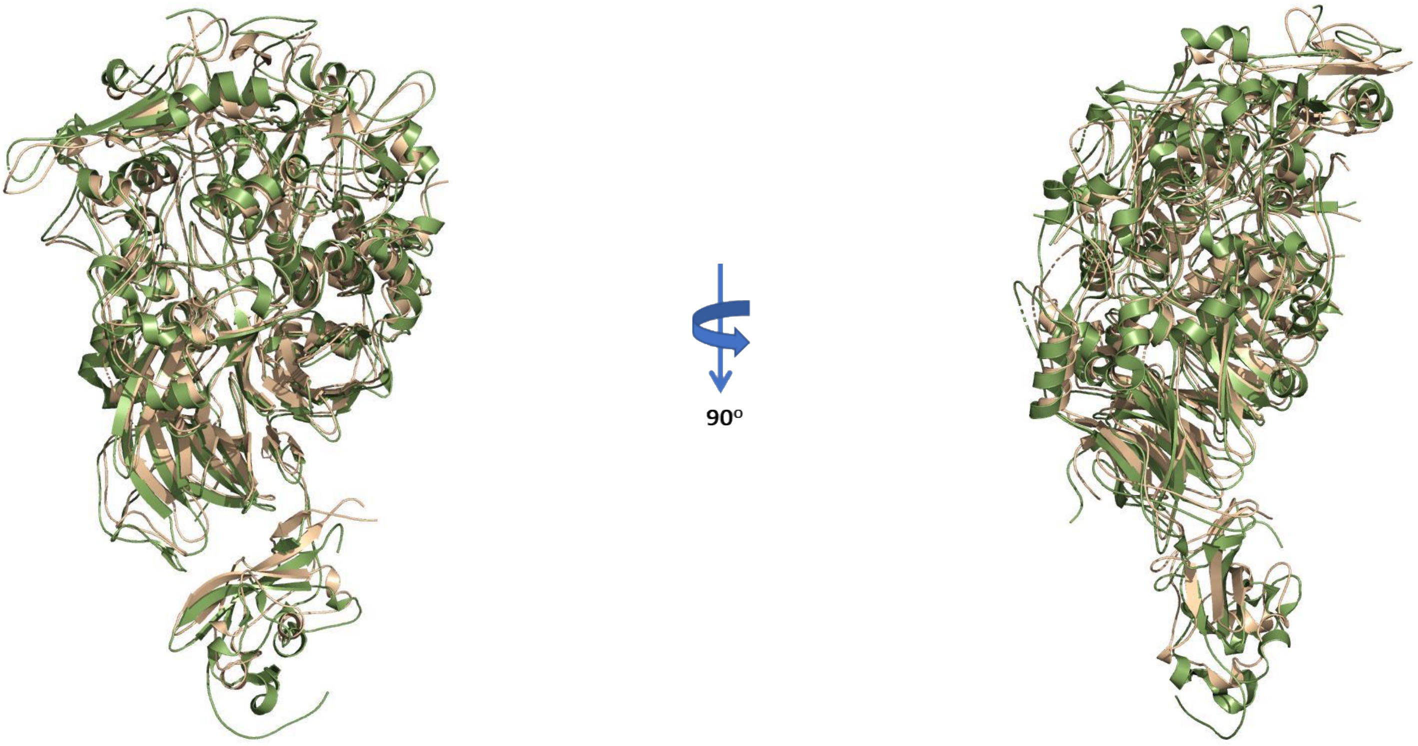


**Supplementary Figure 7. Superposition of P1 structures.** Two 90° apart views of the superposition of the P1 structures determined by X-ray crystallography (PDB code 6RC9) and in this work in the complex P1-Fab(P1/MCA4) (in brown and green, respectively).

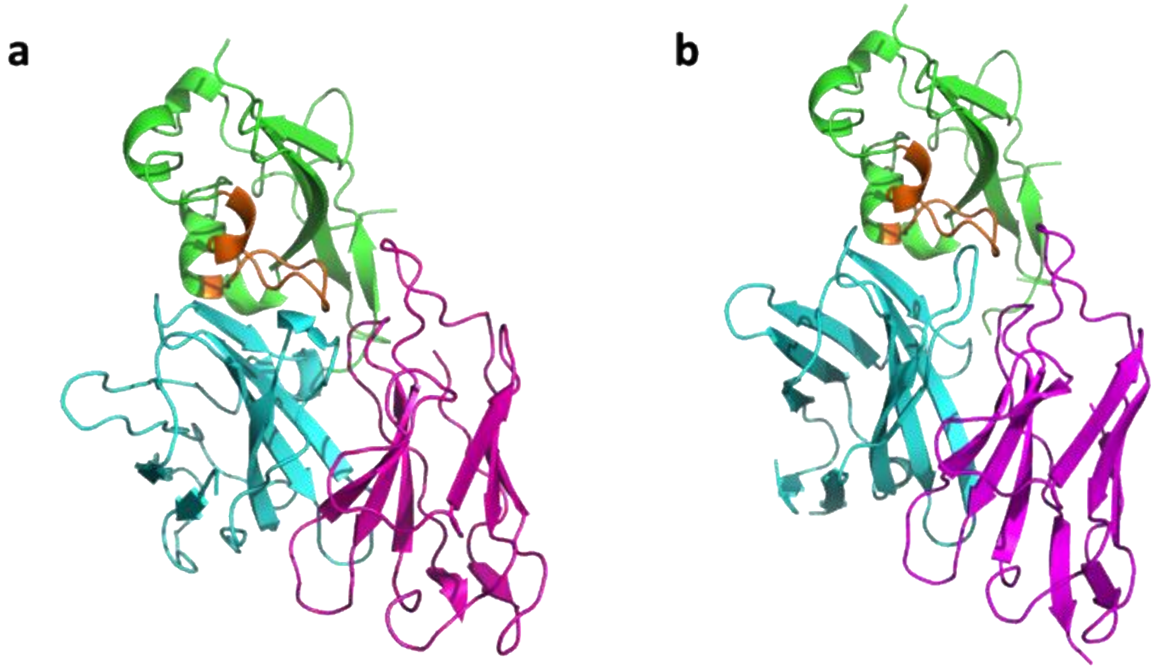


**Supplementary Figure 8. Docking calculations on the binding interface between the Fab and the C-terminal domain of P1. a)** Interface observed in the 2.3 Å resolution map. (b) Top-ranked model obtained from HADDOCK[1] docking simulations performed with a lower resolution map, the P1 crystal structure and the AlphaFold2[2] prediction of the Fab. Docking models were ranked based on Haddock docking score (-66.44 arbitrary units), buried surface area (1457 Å$^2$), and correlation with the cryo-EM map (0.85, as calculated with Chimera[3]). The C-terminal domain of P1 is colored in green, the loop Val1425-Asp1438 in orange, and the heavy and light chains of the Fab in cyan and magenta, respectively. For clarity, only the variable fragment of the Fab is shown.

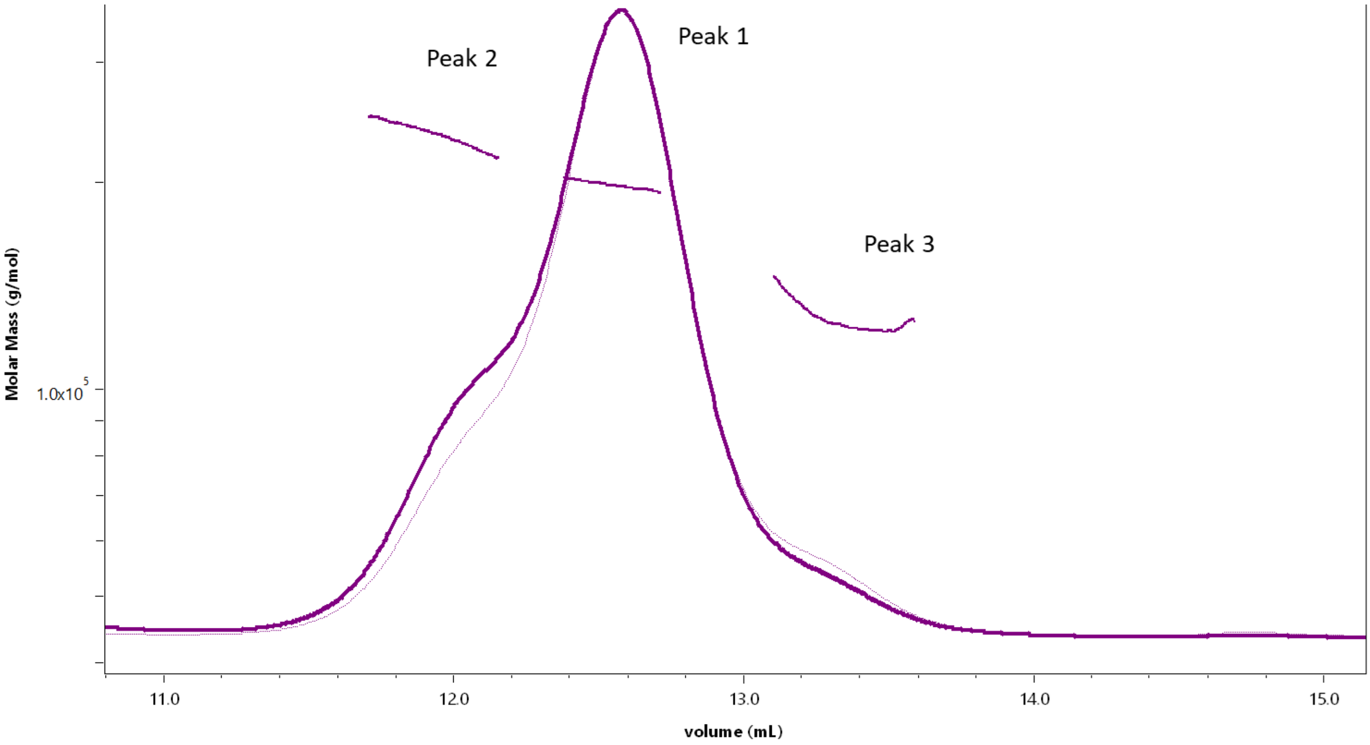


| Peak | Mw (KDa) | Mass Fraction (%) |
|---|---|---|
| Peak 1 | 198.48±0.12 | 62.60 |
| Peak 2 | 231.69±0.29 | 23.55 |
| Peak 3 | 129.04±0.65 | 13.84 |

**Supplementary Figure 9**. **MALS analysis of a sample containing P1, P40/P90 and Fab(P1/MCA4).** Analysis by MALS of a purified sample containing equimolecular amounts of P1, P40/P90 and the Fab(P1/MCA4), indicated the presence of a mixture of species, although none corresponding clearly to the ternary complex.

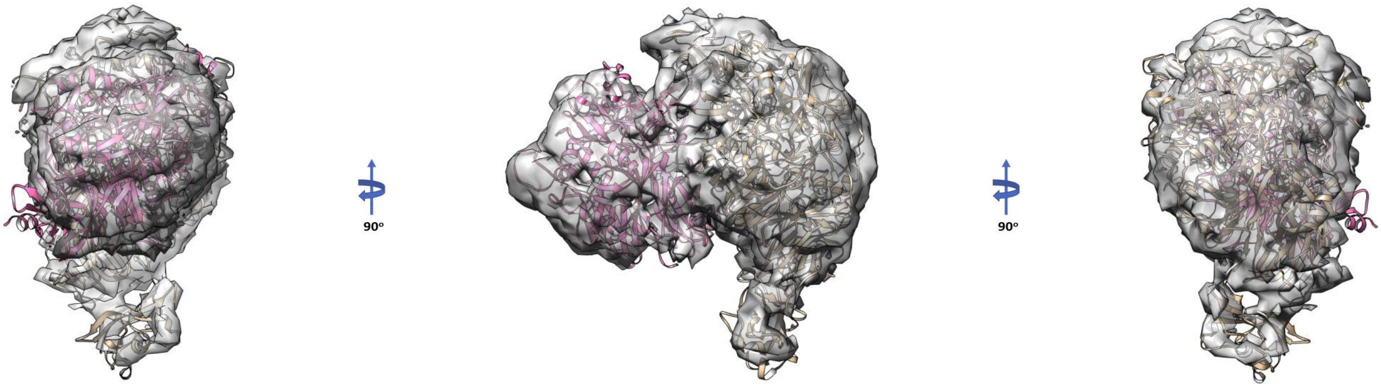


**Supplementary Figure 10**. **Cryo-EM structure determination of the ternary P40/P90-P1-Fab(P1/MCA4) complex**

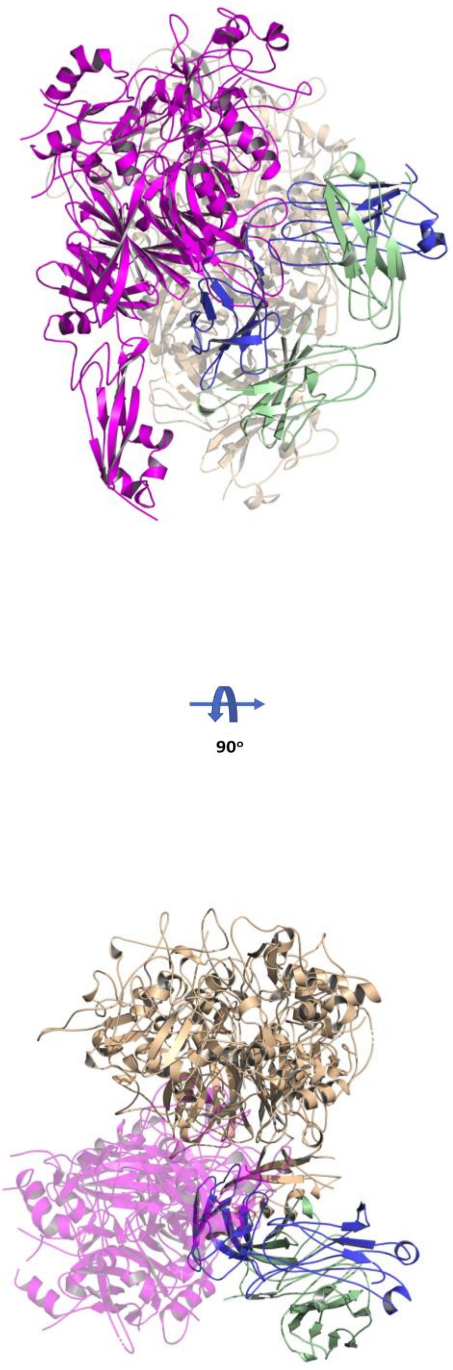


**Supplementary Figure 11. Modelling the ternary complex P40/P90-P1-Fab(P1/MCA4).** Ribbon representation of the ternary complex modelled using the information obtained in the cryo-EM attempts. Steric clashes between P40/P90 (cyan) and the Fab light chain (blue) (heavy chain in green) when they are bound to P1 (brown) can explain the difficulties to obtain the ternary complex.

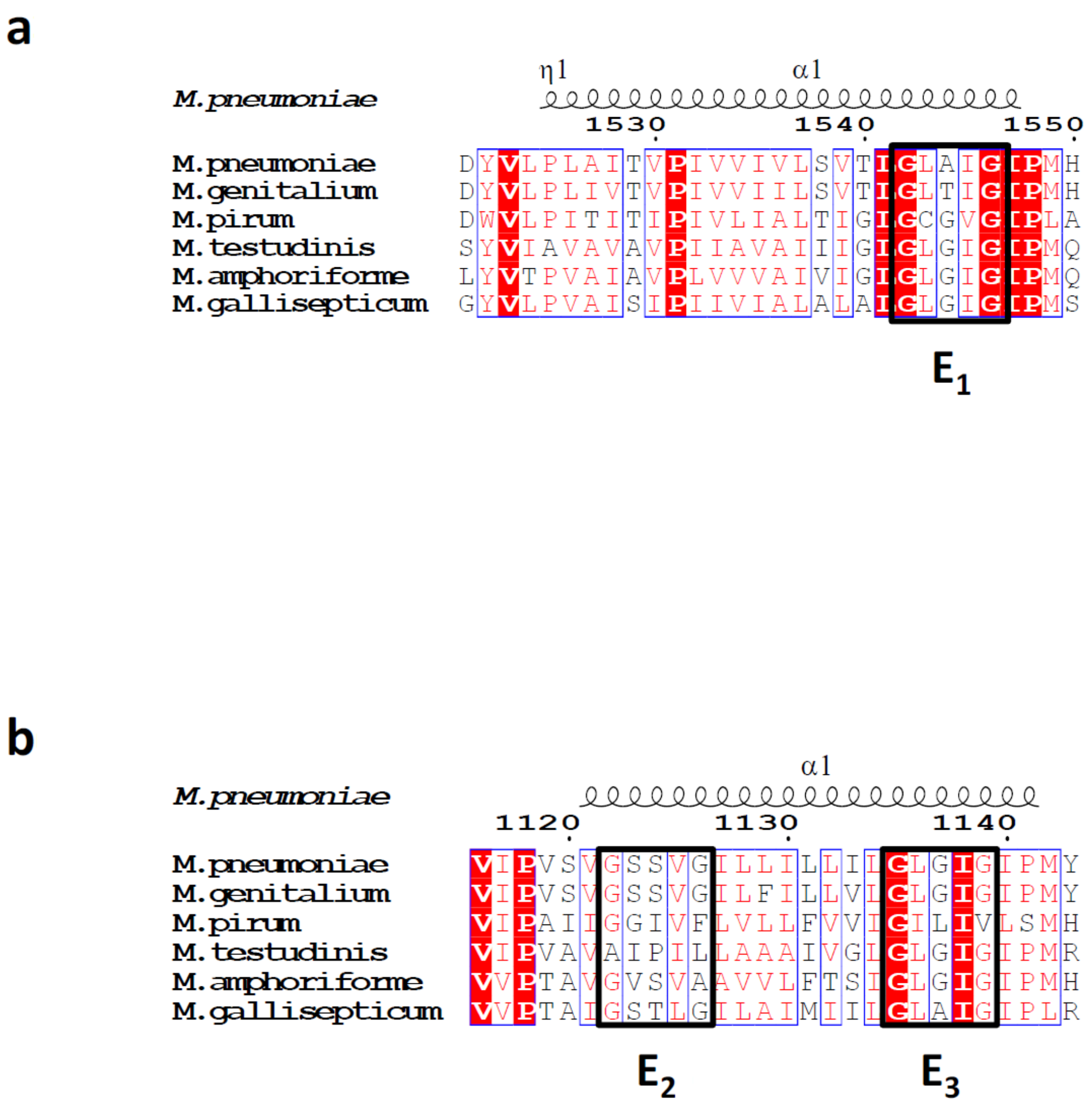


**Supplementary Figure 12. Sequence alignments of transmembrane helices from the adhesins of species belonging to the pneumonia cluster of mycoplasmas.** Orthologues from P1 **(a)** and from P40/P90 **(b).**

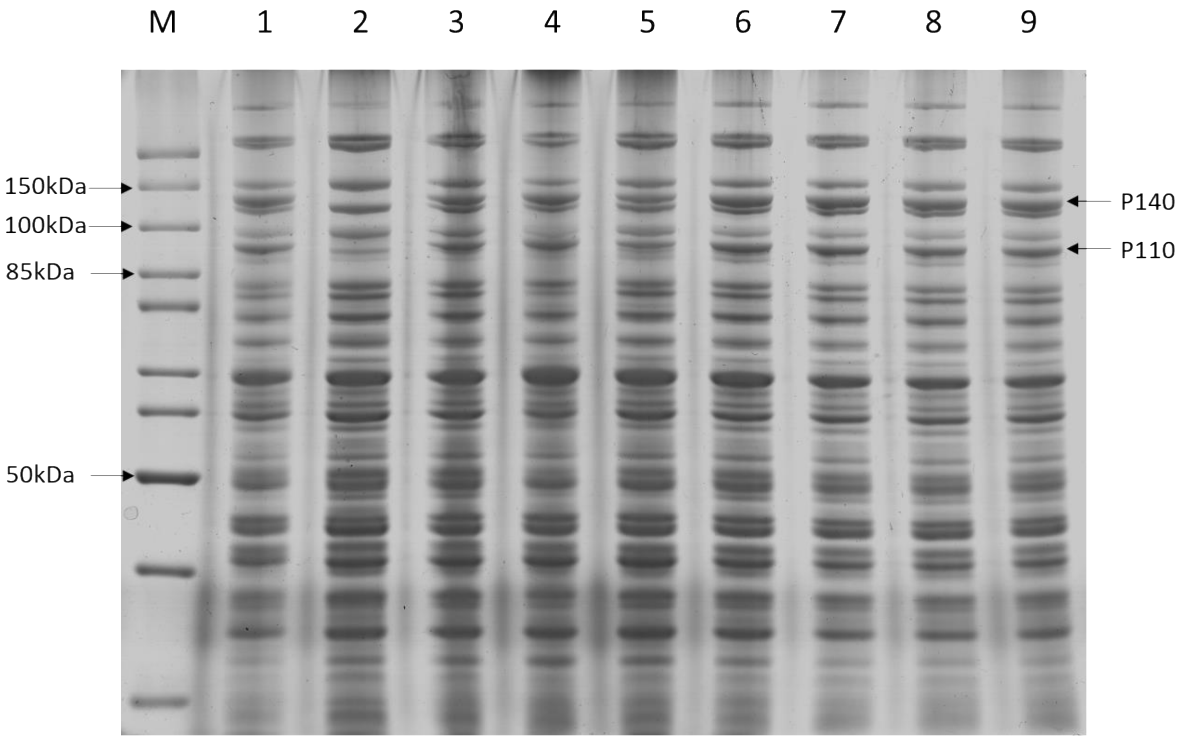


**E1: G1372F-G1376F**

1371Leu-**Gly**-Leu-Thr-Ile-**Gly**-Ile → 1371Leu-**Phe**-Leu-Thr-Ile-**Phe**-Ile

225680-TTG**GGA**TTAACGATT**GGA**ATT → 225680-TTG**TTT**TTAACGATT**TTT**ATT

**E2: G947F-G951F**

946Val-**Gly**-Ser-Ser-Val-**Gly**-Ile → 946Val-**Phe**-Ser-Ser-Val-**Phe**-Ile

228741-GTA**GGT**TCTTCAGTT**GGG**ATC → 228741-GTA**TTT**TCTTCAGTT**TTT**ATC

**E3: G960F-G964F**

959-Leu-**Gly**-Leu-Gly-Ile-**Gly**-Ile → 959-Leu-**Phe**-Leu-Gly-Ile-**Phe**-Ile

228780-TTA**GGA**CTTGGGATT**GGG**ATC → 228780-TTA**TTT**CTTGGGATT**TTT**ATC

**Supplementary Figure 13. SDS-PAGE with protein extracts from *M. genitalium* mutant strains.** 1) *Mge*-WT (G37); 2) G37 ΔAdh; 3) G37 ΔAdh::COM (E1;E2;E3); 4) G37 ΔAdh::COM (E1); 5) G37 ΔAdh::COM (MutE2;MutE3); 6) G37 ΔAdh::COM (E2); 7) G37 ΔAdh::COM (E3); 8) G37 ΔAdh::COM (E1;E2); 9) G37 ΔAdh::COM (E1;E3)

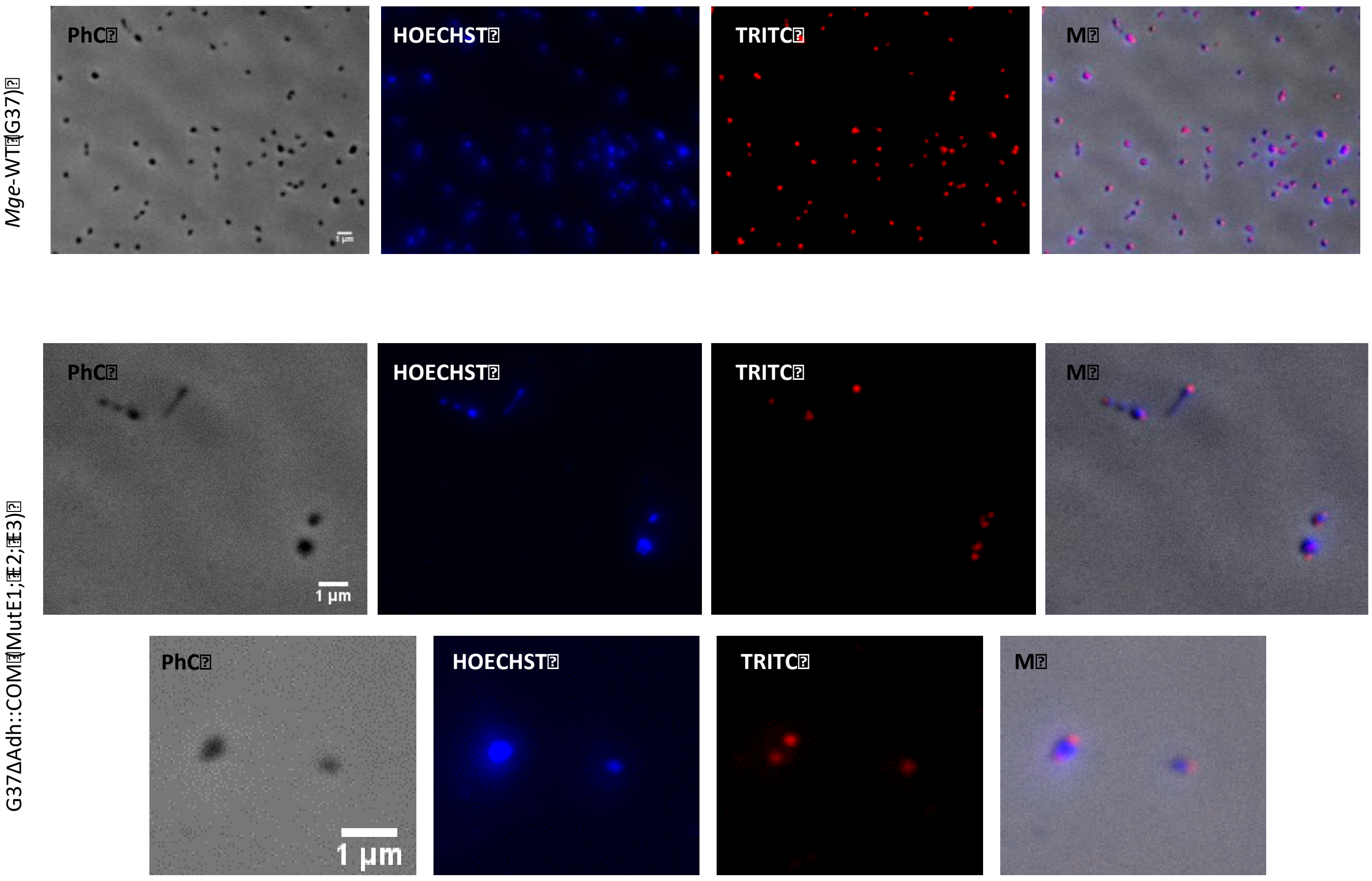


**Supplementary Figure 14. Immunolabeling for the localization of the GPCA complexes in *M. genitalium* WT and MutE1; E2; E3 cells.** Phase contrast (PhC) immunofluorescence microscopy images of cells using labeling with polyclonal antibodies against GPCAs. Labelling for the GPCA seems to concentrates at the tips of the cells.

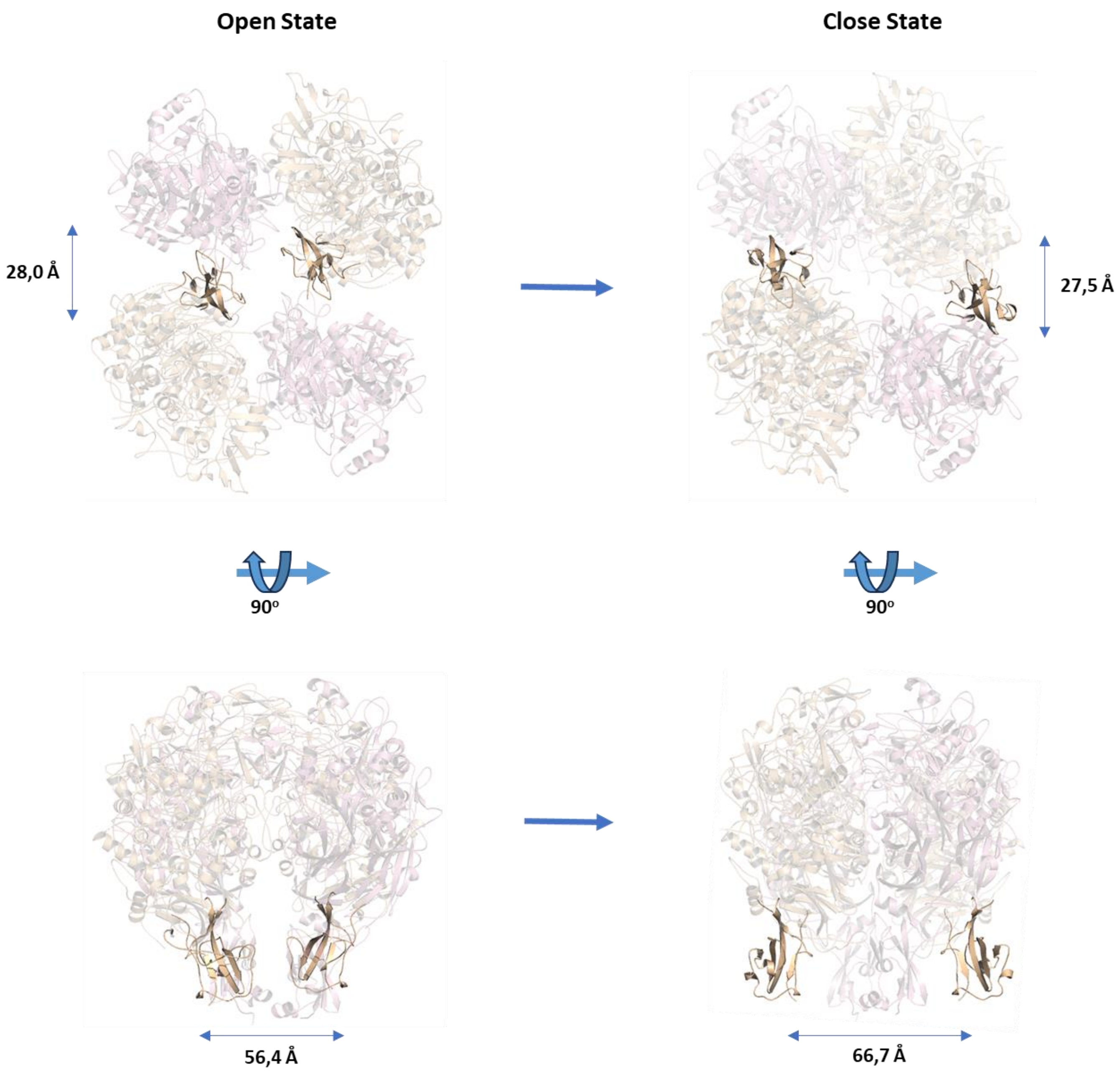


**Supplementary Figure 15. C-terminal domain of P1 movements during the "open" to "closed" transition.** Top and side views (upper and lower panels, respectively) of the GPCA ectodomains in the "open" and "closed" conformations (left and right panels, respectively). Values correspond to the distances (in that perspective) between the C-ends of the C-terminal domains of the two P1 subunits in the GPCA. Besides the linear displacements the C-terminal domain experiences a hinge rotation of about 175º.
