## Supplementary material for "Dynamics of the adhesion complex of the human pathogens *Mycoplasma pneumoniae and Mycoplasma genitalium*": Video legends

### Legends to supplementary videos

**Video I. Microcinematography of *M. pneumoniae* cells in the presence of P1/MCA4 monoclonal antibody.** The clock counter timer at the top-left side of the movie shows the actual microcinematography time in mm:ss format (m: minutes; s: seconds). P1/MCA4 at a final concentration of 10 µg $mL^{-1}$ was added to the culture medium at minute 1:10. This microcinematography was performed with no gelatin added to the culture medium.

**Video II. Microcinematography of *M. pneumoniae* cells in the presence of P1 ectodomain polyclonal antisera.** Mycoplasma cells were in the presence of SP4 medium supplemented with 3 % gelatin. Frames were taken each 0.5 seconds of observation and are showed at 20 frames per second in the movie. The blank frame/s in the first seconds of the movie denote when the polyclonal antisera was added to the cells.

**Video III. Microcinematography of *M. pneumoniae* cells in in the presence P1 N-term polyclonal antisera.** Mycoplasma cells were in the presence of SP4 medium supplemented with 3 % gelatin. Frames were taken each 0.5 seconds of observation and are showed at 20 frames per second in the movie. The blank frame/s in the first seconds of the movie denote when the polyclonal antisera was added to the cells.

**Video IV. Microcinematography of *M. pneumoniae* cells in in the presence of P40/P90 ectodomain polyclonal antisera.** Mycoplasma cells were in the presence of SP4 medium supplemented with 3 % gelatin. Frames were taken each 0.5 seconds of observation and are showed at 20 frames per second in the movie. The blank frame/s in the first seconds of the movie denote when the polyclonal antisera was added to the cells.

**Video V. Modeling the GPCA attachment and detachment cycle and halting of the cycle by the interaction with a Fab**. The video first shows the side and top views of the GPCA complex transitioning from an open conformation (where sialic acid binding is possible) to a closed conformation (sialic acid release). In this transition, the displacement of the transmembrane helix is linked to the hinge rotation of the C-terminal domain from P1. Moreover, the N-terminal domain also moves interacting tightly with the N-domain of P40/P90, which forces the release of the sialic acid from its binding site. This attachment and detachment cycle is interrupted when a Fab interacts with P1 and traps the closed conformation of the C-domain from P1, impeding the complex to reach again the open conformation.
