## Supplementary tables for "Dynamics of the adhesion complex of the human pathogens *Mycoplasma pneumoniae and Mycoplasma genitalium*"

#### Supplementary Table I
#### MCA Monoclonal antibodies generated against the rP1 antigen (A1160-Q1518)

| Clone No. | ELISA screening against rP1 | Western blot against M129 lysate | Binding to live M129 cells | Inhibition of hemadsorption (HA) | Inhibition of gliding |
|---|---|---|---|---|---|
| 3 | + | + | + | + | + |
| 4 | + | + | + | + | + |
| 5 | + | + | + | + | + |
| 8 | + | + | + | + | + |
| 18 | + | + | + | + | + |
| 102 | + | - | - | - | - |
| 104 | + | + | - | - | - |
| 115 | + | + | - | - | - |
| 127 | + | + | - | - | - |
| 128 | + | + | - | - | - |

**Supplementary Table II**

**Cryo-EM data collection of the P1-Fab and of the P1-P40/P90-(Fab) complexes and model refinement of the P1-Fab complex**

| | **P1-Fab** | **P40/P90-P1** |
|---|---|---|
| **PDB ID** | 8ROR | - |
| **Data collection** | | |
| Microscope | CRYO ARM 300 (JEOL) | CRYO ARM 300 (JEOL) |
| Voltage (kV) | 300 | 300 |
| Detector | K3 direct (Gatan) | K3 direct (Gatan) |
| Data collection software | SerialEM | SerialEM |
| Energy Filter | Slit width of 20 eV | Slit width of 20 eV |
| Electron dose ($e^-$/ Å$^2$)/Frame | 2.0 | 2.0 |
| Pixel size (Å) | 0.49 | 0.87 |
| Defocus range (μm) | -0.5 to -1.5 | -0.8 to -1.8 |
| Frames | 50 | 40 |
| Movie number | 27122 | 7983 |
| **Data processing** | | |
| Processing software | RELION-3.1 | cryoSPARC2 |
| Number of extracted particles | 4312408 | 9565198 |
| EMDB code | EMD-19402 | EMD-19403 |
| Map resolution (Å) at: FSC=0.143 (unmasked/masked) | 2.39 | <10 |
| **Refinement** | | |
| Software | PHENIX | |
| Models used (PDB code) | 6RC9 | |
| Atoms (Non-H) | 14210 | |
| Protein residues | 1789 | |
| Ligands | - | |
| Waters | 269 | |
| Bonds (r.m.s.d) | | |
| Length (Å) | 0.004 | |
| Angles (o) | 0.659 | |
| B-factor (Å$^2$) | 83.53 | |
| MolProbity score | 2.72 | |
| Clashcore | 27.36 | |
| Rotamer outliers (%) | 3.79 | |
| Ramachandran plot | | |
| Favored (%) | 94.82 | |
| Allowed (%) | 5.07 | |
| Disallowed (%) | 0.11 | |
| CC (volume) | 0.83 | |
| CC (mask) | 0.83 | |

### Supplementary Table III
### Monoclonal/polyclonal antibodies inhibition assays (extended data)

| | | Before AS | | | | After AS | | | | | | |
|---|---|---|---|---|---|---|---|---|---|---|---|---|
| **PCA** | **MCN replicate** | **Total** | **Non-Motile** | **Motile** | **%Motile Cells** | **Total** | **Non-Motile** | **Motile** | **%Motile Cells** | **$RT_{50}$ (min)** | **Detached cells** | **% Detached cells** |
| **P1** | **1** | 436 | 44 | 392 | 89,9 | 111 | 105 | 6 | 5,4 | 6,95 | 325 | 74,5 |
| | **2** | 225 | 22 | 203 | 90,2 | 73 | 73 | 0 | 0,0 | 3,99 | 152 | 67,6 |
| | **3** | 135 | 16 | 119 | 88,1 | 34 | 34 | 0 | 0,0 | 3,00 | 101 | 74,8 |
| | | | | **Mean** | **89,4** | | | **Mean** | **1,8** | **4,65** | | **72,3** |
| | | | | **SE** | **0,6** | | | **SE** | **1,8** | **1,19** | | **2,4** |
| **P40/P90** | **1** | 464 | 40 | 424 | 91,4 | 133 | 40 | 93 | 69,9 | NA | 331 | 71,3 |
| | **2** | 110 | 18 | 92 | 83,6 | 89 | 19 | 70 | 78,7 | NA | 21 | 19,1 |
| | | | | **Mean** | **87,5** | | | **Mean** | **74,3** | | | **45,2** |
| | | | | **SE** | **3,9** | | | **SE** | **4,4** | | | **26,1** |
| **P1N-ter** | 1 | 575 | 25 | 550 | 95,7 | 231 | 30 | 201 | 87,0 | NA | 344 | 59,8 |
| | 2 | 581 | 26 | 555 | 95,5 | 234 | 38 | 196 | 83,8 | NA | 347 | 59,7 |
| | | | | **Mean** | **95,6** | | | **Mean** | **85,4** | | | **59,8** |
| | | | | **SE** | **0,1** | | | **SE** | **1,6** | | | **0,1** |
| **Negative Control** | **1** | 242 | 21 | 221 | 91,3 | 123 | 21 | 102 | 82,9 | NA | 119 | 49,2 |
| | **2** | 110 | 10 | 100 | 90,9 | 60 | 13 | 47 | 78,3 | NA | 50 | 45,5 |
| | **3** | 302 | 38 | 264 | 87,4 | 121 | 40 | 81 | 66,9 | NA | 181 | 59,9 |
| | | | | **Mean** | **89,9** | | | **Mean** | **76,1** | | | **51,5** |
| | | | | **SE** | **1,2** | | | **SE** | **4,8** | | | **4,3** |

| | | Before AS | | | | After AS | | | | |
|---|---|---|---|---|---|---|---|---|---|---|
| **PCA** | **MCN replicate** | **Total** | **Non-Motile** | **Motile** | **%Motility** | **Total** | **Non-Motile** | **Motile** | **%Motility** | **$RT_{50}$ (min)** |
| **P1** | **1** | 175 | 5 | 170 | 97,1 | 172 | 172 | 0 | 0 | 3,40 |
| | **2** | 162 | 6 | 156 | 96,3 | 168 | 168 | 0 | 0 | 3,04 |
| | **3** | 415 | 11 | 404 | 97,3 | 408 | 338 | 70 | 17,2 | 5,38 |
| | **4** | 360 | 8 | 352 | 97,8 | 372 | 361 | 11 | 3,0 | 4,61 |
| | **5** | 527 | 20 | 507 | 96,2 | 424 | 424 | 0 | 0 | 4,08 |
| | **6** | 692 | 40 | 652 | 94,2 | 692 | 692 | 0 | 0 | 3,79 |
| | | | | **Mean** | **96,5** | | | **Mean** | **3,4** | **4,05** |
| | | | | **SE** | **0,5** | | | **SE** | **2,8** | **0,35** |
| **P40/P90** | **1** | 181 | 7 | 174 | 96,1 | 149 | 7 | 142 | 95,3 | NA |
| | **2** | 313 | 9 | 304 | 97,1 | 340 | 13 | 327 | 96,2 | NA |
| | **3** | 216 | 9 | 207 | 95,8 | 183 | 8 | 175 | 95,6 | NA |
| | | | | **Mean** | **96,4** | | | **Mean** | **95,7** | |
| | | | | **SE** | **0,4** | | | **SE** | **0,3** | |
| **P1N-ter** | **1** | 524 | 52 | 472 | 90,1 | 441 | 55 | 386 | 87,5 | NA |
| | **2** | 424 | 87 | 337 | 79,5 | 453 | 87 | 366 | 80,8 | NA |
| | **3** | 747 | 49 | 698 | 93,4 | 747 | 54 | 693 | 92,8 | NA |
| | **4** | 588 | 43 | 545 | 92,7 | 588 | 49 | 539 | 91,7 | NA |
| | | | | **Mean** | **88,9** | | | **Mean** | **88,2** | |
| | | | | **SE** | **3,2** | | | **SE** | **2,7** | |
| **Negative Control** | **1** | 117 | 6 | 111 | 94,9 | 114 | 6 | 108 | 94,7 | NA |
| | **2** | 230 | 8 | 222 | 96,5 | 257 | 12 | 245 | 95,3 | NA |
| | **3** | 176 | 6 | 170 | 96,6 | 177 | 6 | 171 | 96,6 | NA |
| | **4** | 124 | 4 | 120 | 96,8 | 125 | 4 | 121 | 96,8 | NA |
| | **5** | 417 | 18 | 399 | 95,7 | 418 | 16 | 402 | 96,2 | NA |
| | | | | **Mean** | **96,3** | | | **Mean** | **96,5** | |
| | | | | **SE** | **0,3** | | | **SE** | **0,1** | |

| | | Before AS | | | | After AS | | | | |
|---|---|---|---|---|---|---|---|---|---|---|
| **MCA** | **MCN replicate** | **Total** | **Non-Motile** | **Motile** | **%Motility** | **Total** | **Non-Motile** | **Motile** | **%Motility** | **$RT_{50}$ (min)** |
| **P1-MCA4** | **1** | 349 | 5 | 344 | 98,6 | 349 | 349 | 0 | 0 | 2,63 |
| | **2** | 184 | 6 | 178 | 96,7 | 168 | 168 | 0 | 0 | 2,18 |
| | **3** | 142 | 5 | 137 | 96,5 | 142 | 142 | 0 | 0 | 3,09 |
| | | | | **Mean** | **97,3** | | | **Mean** | **0** | **2,6** |
| | | | | **SE** | **0,7** | | | **SE** | **0** | **0,26** |

**Supplementary Table IV**
**Engelman mutants**

| Strain | Synonyms | Mutation Description |
|---|---|---|
| **G37** | *Mge*-WT | - |
| **G37ΔAdh** | G37ΔMG_191-ΔMG_192 | Deletion of the MG_191 and MG_192 genes by allelic exchange. |
| **G37ΔAdh::COM** | G37ΔMG_191-ΔMG_192::MG_191-MG_192 | Re-introduction, by transposon insertion, of a MG_191 and MG_192 wild-type alleles in a G37ΔAdh mutant. It is resistant to puromycin. |
| **G37ΔAdh::COM (MutE1)** | G37ΔAdh::COM (E1) | Re-introduction of a MG_191 and MG_192 alleles bearing **P140: G1372F-G1376F** substitutions, in a G37ΔAdh mutant. It is resistant to puromycin. |
| **G37ΔAdh::COM (MutE2; MutE3)** | G37ΔAdh::COM (E2; E3) | Re-introduction of a MG_191 and MG_192 alleles bearing **P110a: G947F-G951F** and **P110b: G960F-G964F** substitutions, in a G37ΔAdh mutant. It is resistant to puromycin. |
| **G37ΔAdh::COM (MutE1; MutE2; MutE3)** | G37ΔAdh::COM (E1; E2; E3) | Re-introduction of a MG_191 and MG_192 alleles bearing **P140: G1372F-G1376F**, **P110a: G947F-G951F** and **P110b: G960F-G964F** substitutions, in a G37ΔAdh mutant. It is resistant to puromycin. |
| **G37ΔAdh::COM (MutE2)** | G37ΔAdh::COM (E2) | Re-introduction of a MG_191 and MG_192 alleles bearing **P110a: G947F-G951F** substitutions, in a G37ΔAdh mutant. It is resistant to puromycin. |

| Strain | Synonyms | Mutation Description |
|---|---|---|
| **G37ΔAdh::COM (MutE3)** | G37ΔAdh::COM (E3) | Re-introduction of a MG_191 and MG_192 alleles bearing **P110b: G960F-G964F** substitutions, in a G37ΔAdh mutant. It is resistant to puromycin. |
| **G37ΔAdh::COM (MutE1; MutE2)** | G37ΔAdh::COM (E1; E2) | Re-introduction of a MG_191 and MG_192 alleles bearing **P140: G1372F-G1376F** and **P110a: G947F-G951F** substitutions, in a G37ΔAdh mutant. It is resistant to puromycin. |
| **G37ΔAdh-COM (MutE1; MutE3)** | G37ΔAdh-COM (E1; E3) | Re-introduction of a MG_191 and MG_192 alleles bearing **P140: G1372F-G1376F** and **P110b: G960F-G964F** substitutions, in a G37ΔAdh mutant. It is resistant to puromycin. |

**Supplementary Table V**
**Primers used for P1 and P40/P90 protein constructs expression**

| Name | Sequence (5'→3') |
|---|---|
| P1F | AGGAGATATACCATGACCGTGGTTGGTCACTTTACC |
| P1R | GTGATGGTGATGTTTATCCGGCCACTGGTTGAACGG |
| P1F_2 | AGGCCATGGCGGCCTTTCGTGGCAGTTG |
| P1R_2 | GTGCTCGAGTCATAAATACTAAGCGGGTT |
| P1Ct1400_F | AGGAGATATACCATGGCGGATACCGGTCCGCAG |
| P1Ct1376_F | AGGAGATATACCATGAAAATGAACGATGACGTTG |
| P40P90_F | AGGAGATATACCATGAGCCTGGCGAACACCTATCTGCTG |
| P40P90_R | GTGATGTGTATGTTTGCTCGGCACGCGCCGCAAAACC |

#### Supplementary Table VI
#### Plasmids used for expression of P1 protein fragments for epitope mapping of P1/MCA4

| No. | Plasmids | Primers used for plasmid construction* | | Plasmid size | Recombinant P1 fragment | M.W. (kDa) ** | Bind to P1/MC4*** |
|---|---|---|---|---|---|---|---|
| 1 | pP1-8 | 5'-AGGCCATGGCGGCCTTTCGTGGCAGTTG -3' | 5'-GTGCTCGAGTCATAAATACTAAGCGGGTT -3' | 6789 | A1160 - Q1518 | 43,4 | + |
| 2 | pP1-8d-N1 | 5'-TTATTGATTAGTATTTATGACTCGAGCAC -3' | 5'-AATACTAATCAATAAACAGCGGTATGT -3' | 6246 | A1160 - D1446 | 35,4 | + |
| 3 | pP1-8d-N2 | 5'-CGCTGTTTTAGTATTTATGACTCGAGCAC -3' | 5'-AATACTAAAACAGCGGTATGTACTGGT -3' | 6240 | A1160 - F1444 | 35,2 | + |
| 4 | pP1-8d-N3 | 5'-ACATACCGTAGTATTTATGACTCGAGCAC -3' | 5'-AATACTACGGTATGTACTGGTCATACA -3' | 6234 | A1160 - P1442 | 34,9 | + |
| 5 | pP1-8d-N4 | 5'-ACCAGTACTAGTATTTATGACTCGAGCAC -3' | 5'-AATACTAGTACTGGTCATACACCAACA -3' | 6228 | A1160 - Y1440 | 34,7 | + |
| 6 | pP1-8d-N5 | 5'-TGTATGACTAGTATTTATGACTCGAGCAC -3' | 5'-AATACTAGTCATACACCAACATAGTTA -3' | 6222 | A1160 - D1438 | 34,4 | + |
| 7 | pP1-8d-N6 | 5'-TGGTGTATTAGTATTTATGACTCGAGCAC -3' | 5'-AATACTAATACACCAACATAGTTACCG -3' | 6219 | A1160 - Y1437 | 34,3 | - |
| 8 | pP1-8d-N7 | 5'-TGTTGGTGTAGTATTTATGACTCGAGCAC -3' | 5'-AATACTACACCAACATAGTTACCGGAT -3' | 6216 | A1160 - V1436 | 34,1 | - |
| 9 | pP1-8d-N8 | 5'-TAACTATGTAGTATTTATGACTCGAGCAC -3' | 5'-AATACTACATAGTTACCGGATCAAACA -3' | 6210 | A1160 - M1434 | 33,9 | - |

| No. | Plasmids | Primers used for plasmid construction* | | Plasmid size | Recombinant P1 fragment | M.W. (kDa) ** | Bind to P1/MC4 |
|---|---|---|---|---|---|---|---|
| 10 | pP1-8d-N9 | 5'-ATCCGGTATAGTATTTATGACTCGAGCAC -3' | 5'-AATACTATACCGGATCAAACAGATCGG -3' | 6204 | A1160 – V1432 | 33,7 | - |
| 11 | pP1-8d-C1 | 5'-AGGCCATGACTATGTTGGTGTATGACCA -3' | 5'-ACATAGTCATGGCCTTGTCGTCGTC -3' | 5970 | T1433 - Q1518 | 14,7 | - |
| 12 | pP1-8d-C2 | 5'-AGGCCATGCCGGTAACTATGTTGGTGTA -3' | 5'-TTACCGGCATGGCCTTGTCGTCGTC -3' | 5976 | P1431 - Q1518 | 14,9 | - |
| 13 | pP1-8d-C3 | 5'-AGGCCATGTTTGATCCGGTAACTATGTT -3' | 5'-GATCAAACATGGCCTTGTCGTCGTC -3' | 5982 | F1429 - Q1518 | 15,2 | - |
| 14 | pP1-8d-C4 | 5'-AGGCCATGGATCTGTTTGATCCGGTAAC -3' | 5'-ACAGATCCATGGCCTTGTCGTCGTC -3' | 5988 | D1427 - Q1518 | 15,4 | - |
| 15 | pP1-8d-C5 | 5'-AGGCCATGACCGATCTGTTTGATCCGGT -3' | 5'-GATCGGTCATGGCCTTGTCGTCGTC -3' | 5991 | T1426 - Q1518 | 15,5 | + |
| 16 | pP1-8d-C6 | 5'-AGGCCATGGTCACCGATCTGTTTGATCC -3' | 5'-CGGTGACCATGGCCTTGTCGTCGTC -3' | 5994 | V1425 - Q1518 | 15,6 | + |
| 17 | pP1-8d-C7 | 5'-AGGCCATGGCTGACACTGGTCCACAA -3' | 5'-TGTCAGCCATGGCCTTGTCGTCGTC -3' | 6069 | A1400 – Q1518 | 18,4 | + |
| 18 | pP1-8d-C8 | 5'-AGGCCATGAAGATGAATGACGATGTT -3' | 5'-TCATCTTCATGGCCTTGTCGTCGTC -3' | 6141 | K1376 – Q1518 | 21 | + |

The pP1-8 plasmid was constructed by inserting a PCR amplified fragment of p1 gene into the NcoI and XhoI sites of pET-30c(+) expression vector.

The remaining plasmids were constructed by deletion of the pP1-8 using specific primer set and PrimeSTAR Mutagenesis Basal kit.

* NcoI and XhoI sites of the primers used for pP1-8 plasmid construction are underlined.

** Molecular weight of recombinant P1 fragments including His- and S-tag sequences (45 aa: 5 kDa) from the pET-30c(+) expression vector.

### Supplementary Table VII
### Primers used in the generation of M. genitalium adhesins constructs

| Primer Name | Sequence 5' -3' | Use |
|---|---|---|
| | **Double adhesins null mutant** | |
| MgParUp-F | ATCATTACCATTATCAATG | Build pBEΔAdh |
| MgParUp-R | GGATCCATGCACCTCTCGAGACAAACTTAATTATAAACAAT | |
| MgParDw-F | CTCGAGAGGTGCATGGATCCTAGTTTTTAACCTTTCAATAAC | |
| MgParDw-R | CTACTATTGCTAGGTTTCAC | |
| Lox71p438Fwd-XhoI | CTC GAG TAC CGT TCG TAT AAT GTA TGC TAT ACG AAG TTA TTA GTA TTT AGA ATT AAT AAA GT | |
| CatLox66Rev-BamHI | GGA TCC TAC CGT TCG TAT AGC ATA CAT TAT ACG AAG TTA TTT ACG CCC CGC CCT GCC ACT | |
| | **Complementation of the null mutant** | |
| COMmg191-F-ApaI | AGTGGGCCCACTAACAAAAAACAAATTAGTGATG | Build pMTnPacCOM (MG_191/MG_192) |
| COMmg191/192-R-SalI | AGTGTCGACATCCACTCTCTAAATTGCAAGTTTAG | |
| | **Engelman Motif mutants** | |
| EngMotifP140-F | TTTTTAACGATTTTTATTCCAATGCACAGAAAC | Build pMTnPacCOM:MG_191/MG_192 with the different Engelman Motif substitutions |
| EngMotifP140-R | *CAAAGTAACACTAAGGATAATC | |
| EngMotifP110-F | TTTTCTTCAGTTTTTATCTTGTTTATCTTGTTAGTCTTATTTCTTGGGATTTTTATCCCAATGTACAGGGTAAGA | |
| EngMotifP110a-F | TTTTCTTCAGTTTTTATCTTGTTTATCTTGTTAGTC | |
| EngMotifP110-R | *TACTGATACAGGGATCACCCA | |
| EngMotifP110b-F | TTTCTTGGGATTTTTATCCCAATGTACAGGGTAAGA | |
| EngMotifP110b-R | *TAAGACTAACAAGATAAACAAG | |

| Primer Name | Sequence 5' -3' | Use |
|---|---|---|
| **Sequencing primers** | | |
| Pac-Up | GTAGCTAATCTAACAGTAGG | To sequence DNA inserts cloned in a miniPac Tnp |
| Pac-Dw | GTCCTAGAACTTGGTGTATG | To determine miniTnPac Tnp insertion point |
| COMmg191-R-SalI | AGTGTCGACttattgttttactggaggttt | To sequence DNA inserts cloned in a miniPac Tnp |
| Fup17 | GTAAAACGACGGCCAGT | Universal Forward primer |
| Rup17 | GGAAACAGCTATGACCATG | Universal Reverse primer |
| Tnp3 | CATGATGAATGGATTTATTC | To sequence DNA inserts cloned in a miniPac Tnp |
| RTPCR192-F | GTTGATACACTCACAACTG | To sequence DNA inserts cloned in a miniPac Tnp |
| RTPCR192-R | CTAACTTTTGGTTTCTTCTGAC | To sequence DNA inserts cloned in a miniPac Tnp |
| RTPCRmg191-F | CTGGAGAGAACCCAGGATCA | To sequence DNA inserts cloned in a miniPac Tnp |
| SEQmg191a-F | GGTTAGTTTCTATGATGCAC | To sequence DNA inserts cloned in a miniPac Tnp |
| SEQmg191b-F | GATACAGCTACTGTACCTAG | To sequence DNA inserts cloned in a miniPac Tnp |
| SEQmg191c-F | GTTCTACCTTCGATCAGTTC | To sequence DNA inserts cloned in a miniPac Tnp |
| SEQmg191d-F | GCATTACTCCATACCTATGG | To sequence DNA inserts cloned in a miniPac Tnp |
| SEQmg191e-F | ATTAACACCATCACCACTAC | To sequence DNA inserts cloned in a miniPac Tnp |
| **Oligonucleotides used in mutant screening** | | |
| SCRMgPaFw | GTCTGTTTGCCATCTATGACA | To screen for MG_191/MG_192 null mutants |
| SCRMgPaRev | GAGCCACCTGAAGTGACTT | |

**Supplementary Table VIII**
**Antibodies used in this work**

| Antibody | Host | Dilution | Conjugated enzyme | Source |
|---|---|---|---|---|
| **Anti-mouse IgG (H+L) Alexa Fluor 555** | Goat | 1:250 | | Invitrogen |
| **Polyclonal anti-Mouse IgG (H+L)** | Goat | 1:3000 | HRP | Bio-Rad |
| **Polyclonal anti-MG_191/MG_192 (Nap)** | Mouse | 1:1000 | - | This work |
| **Polyclonal anti-MPN141 (P1)** | Mouse | 1:10/1:1000 | - | This work |
| **Polyclonal anti-MPN142 (P40P90)** | Mouse | 1:10/1:1000 | - | This work |
| **Polyclonal anti-MPN141 N-domain** | Mouse | 1:10/1:1000 | - | This work |
| **Monoclonal anti-MPN141 (P1/MCA4)** | Mouse? | 1:100/1:3000 | - | Seto *et al.* 2005 (unpublished). This work |
